# Comprehensive computational analysis of the molecular mechanism of self-incompatibility in Brassicaceae using improved structure prediction

**DOI:** 10.1101/2023.06.19.545538

**Authors:** Tomoki Sawa, Yoshitaka Moriwaki, Hanting Jiang, Kohji Murase, Seiji Takayama, Kentaro Shimizu, Tohru Terada

## Abstract

Plants employ self-incompatibility (SI) to promote cross-fertilization. In Brassicaceae, this process is regulated by the formation of a complex between the pistil determinant *S* receptor kinase (SRK) and the pollen determinant *S*-locus protein 11 (SP11, also known as *S*-locus cysteine-rich protein, SCR). In our previous study, we used the crystal structures of two eSRK–SP11 complexes in *Brassica rapa S*_8_ and *S*_9_ haplotypes and nine computationally predicted complex models to demonstrate that only the SRK ectodomain (eSRK) and SP11 pairs derived from the same *S* haplotype exhibit high binding free energy. However, predicting the eSRK–SP11 complex structures for the other 100+ *S* haplotypes and genera remains difficult because of SP11 polymorphism in sequence and structure. Although protein structure prediction using AlphaFold2 exhibits considerably high accuracy for most protein monomers and complexes, 46% of the predicted SP11 structures that we tested showed < 75 mean per-residue confidence score (pLDDT). Here, we demonstrate that the use of curated multiple sequence alignment (MSA) for cysteine-rich proteins significantly improved model accuracy for SP11 and eSRK–SP11 complexes. Additionally, we calculated the binding free energies of the predicted eSRK–SP11 complexes using molecular dynamics (MD) simulations and observed that some *Arabidopsis* haplotypes formed a binding mode that was critically different from that of *B. rapa S*_8_ and *S*_9_. Thus, our computational results provide insights into the haplotype-specific eSRK–SP11 binding modes in Brassicaceae at the residue level. The predicted models are freely available at Zenodo, https://doi.org/10.5281/zenodo.8047768.

## INTRODUCTION

Self-incompatibility (SI) is a mechanism that many flowering plants acquire to prevent self-fertilization and promote genetic diversity^*1, 2*^. Pollen hydration, tube germination, and elongation are rejected if the pollen is genetically identical or closely related to the pistils. Over the course of evolution, plants have acquired various molecular mechanisms for SI that are currently being researched in the Brassicaceae^*3-5*^, Solanaceae^*6-9*^, and Papaveraceae^*10-13*^ families among others. In Brassicaceae, SI is sporophytically controlled by the *S*-locus^*3-5*^. The family Brassicaceae contains diverse genera and species for which extensive surveys of the diversity of the *S*-locus have been performed, including *Brassica rapa* (Chinese cabbage, turnip, etc.), *B. oleracea* (cabbage, broccoli, etc.), *B. napus* (rapeseed, etc.), *Raphanus sativus* (radish, etc.), *R. raphanistrum* (wild radish, etc.), *Arabidopsis lyrata*, and *A. halleri*.

*S*-locus glycoprotein (SLG) was the first identified product of the *S*-locus^*14, 15*^. Subsequently, a receptor-type kinase called *S*-receptor kinase (SRK) with an SLG-like extracellular region was identified as the pistil determinant of SI in *B. oleracea* (*Bo*) and *B. rapa* (*Br*)^*16, 17*^. Furthermore, *S*-locus protein 11 (SP11 or *S*-locus cysteine-rich protein (SCR)) has been identified as a determinant present on the pollen side^*18-21*^. SRK, which is located on the plasma membrane of the papilla cells, specifically binds to its cognate SP11 that is released from the pollen surface. This triggers the self-phosphorylation of the kinase domain inside the cell, which results in the rejection of fertilization^*22-26*^. The alleles of *SLG, SRK*, and *SP11* are tightly linked at the *S*-locus and are transmitted to the progeny as a single set called *S* haplotype^*18, 27*^.

The amino acid sequences of SRK and SP11 are significantly different among the *S* haplotypes. The SRK ectodomain (eSRK) shares two lectin domains with an EGF-like domain and a PAN domain among the haplotypes^*28*^ in addition to three hypervariable (HV I–III) regions with notably different sequence compositions^*29, 30*^. In contrast, SP11 is a defensin-like protein that comprises 60–70 amino acids with four or five disulfide bonds and is characterized by numerous insertions, deletions, and variations across haplotypes^*31*^. Based on sequence similarity, the *S* haplotypes are classified into classes I and II in *Brassica* and *Raphanus. Br* class II comprises the *S*_29_, *S*_40_, *S*_44_, and *S*_60_ haplotypes^*32*^. However, the dominant nature of class I pollen phenotype hinders that of class II. Regulation of pollen dominance is achieved through the suppression of SP11 expression by two small RNAs: *SP11 methylation inducer* (*SMI*) and *SMI2*^*33, 34*^.

Recently, two eSRK–SP11 complex crystal structures have been reported. Ma *et al*. determined the *Br S*_9_-eSRK–*S*_9_-SP11 complex structure, in which the two SP11 molecules are tightly bound to the interface of the two eSRKs^*35*^. Subsequently, we presented the crystal structure of the engineered *Br S*_8_-eSRK–*S*_8_-SP11 complex and nine computational complex models of *S*_8_-, *S*_9_-relative and class-II haplotypes using homology modeling and accelerated molecular dynamics (MD) simulations^*36*^. Additionally, we showed that the calculated binding free energies were largely negative for all 11 cognate eSRK–SP11 complexes but not for the 156 non-cognate ones. We have also validated that a point mutation in the *S*_36_-SP11 residue located at the interface, inferred from the *S*_36_-eSRK–*S*_36_-SP11 complex model, disrupted the SI reaction in the pollination bioassay^*36*^. These results indicate that the use of appropriate predicted structures can lead to a better understanding of the self/nonself-discrimination mechanism between eSRK and SP11 without experimental crystallization. However, attaining a comprehensive understanding of the mechanism encompassing all *S* haplotypes remains challenging because the estimated number of haplotypes is > 100 for *Br*^*4*^ and > 50 for *Bo*^*37*^. Additionally, most SRK and SP11 proteins cannot be readily expressed in vitro, making experimental validation highly difficult. Furthermore, obtaining the stable structures for all eSRK–SP11 pairs through accelerated MD simulations is impractical owing to the substantial computational resources required.

AlphaFold2^*38*^ and AlphaFold-Multimer (AF-Multimer)^*39*^ released in 2021, exhibited high accuracy in predicting both monomeric and complex structures. We present a concise overview of how AlphaFold2/AF-Multimer performs structure prediction: First, multiple sequence alignment (MSA) for the input sequence is retrieved from the sequence database using jackHMMER^*40*^ and HHblits^*41*^, and template structures are searched in the Protein Data Bank (PDB) database using HHSearch^*42*^; second, the Evoformer processes the MSA and template structures to extract their converged evolutionary and spatial relationships (denoted as MSA and pair representations in that paper); third, the Structure Module assembles the protein structure based on the representations, and the output structure is reutilized thrice as the input of the Evoformer. Importantly, the accuracy of the predicted model decreases substantially when the median MSA depth is < 30 sequences. This underscores the substantial influence of both the quantity and quality of MSA on their accuracy.

The great success of AlphaFold2 in the field of protein structure prediction is expected to advance our understanding of proteins based on their structures. ColabFold^*43*^, which is a derivative of AlphaFold2, offers structure prediction through a web browser by leveraging the MMSeqs2 webserver^*44, 45*^ to obtain an MSA file of the input sequence. Moreover, starting from July 2022, AlphaFold Protein Structure Database (AlphaFold DB)^*46*^ has made predicted monomeric structures available for nearly all proteins recorded in the UniProt database. Despite the per-residue model confidence (a pLDDT metric)^*38*^ being reliably high for eSRK model structures in AlphaFold DB, the confidence for SP11 models is relatively low. This discrepancy is attributed to the limited number of homologous sequences available for SP11. An additional crucial fact worth mentioning is that the AlphaFold DB does not provide structural information for protein complexes that are crucial for understanding molecular recognition.

In this study, we report more plausible SP11 and eSRK–SP11 model structures using the cognate eSRK and SP11 sequence pairs that were annotated in previous experimental studies to overcome the drawbacks of structure prediction with AlphaFold2. Moreover, we comprehensively explore the molecular recognition mechanism between cognate eSRK and SP11 pairs at the residue level based on predicted complex models. Consequently, the models suggested that the presence of variable regions other than HV I–III, which may contribute to the self/nonself-discrimination, and the eSRK–SP11 binding mode vary significantly depending on the *S* haplotypes.

## METHODS

### Preparation of amino acid sequences

The amino acid sequences were obtained from GenBank for eSRK and SP11 from seven species, namely, *Brassica rapa* (*Br*), *B. oleracea* (*Bo*), *B. napus* (*Bn*), *R. sativus* (*Rs*), *R. raphanistrum* (*Rr*), *A. lyrata* (*Al*), and *A. halleri* (*Ah*) and used for structural prediction. In total, 98 eSRK and SP11 sequence pairs derived from the same *S*-haplotypes were used in this study^*17-19, 21, 30-32, 47-70*^. The sequences and accession IDs are shown in **Supplementary File 1**.

### Structure prediction with ColabFold with specified MSAs

The structures of eSRK and SP11 proteins were predicted using ColabFold^*43*^, which is a derivative of AlphaFold2. During a typical prediction process, ColabFold uses the MMSeqs2 webserver to retrieve an MSA file of the input amino acid sequence ^*44, 45*^. In this study, we obtained MSA files using the UniRef30^*71*^ (uniref30_2202) and BFD/MGnify (bfd_mgy_colabfold) databases. We denoted the retrieved MSA files as “default” MSA for the comparison described in a later section. In the prediction using ColabFold, the recycle number for the prediction was set to three and the template structure was not used. These predictions were performed using LocalColabFold, which is the command line interface of ColabFold (https://github.com/YoshitakaMo/localcolabfold).

### Construction of MSA for ColabFold

In addition to the “default” MSA, four other MSAs were constructed to improve the predicted models. The CysBar^*72*^ Python script was used to construct MSAs that properly expressed the structural features of the SP11 defensin-like domain. This tool facilitates the alignment of structurally homologous cysteine residues within the MSA. Typically, such homologous Cys residues are identified by combining them with known structures of high homology detected using the DALI server^*73, 74*^. However, owing to the large number of structurally unknown and polymorphic SP11s, the disulfide bond pattern of the *S*_8_- or *S*_9_-SP11 crystal structures was used as the standard, and other SP11s were assumed to form similar pairs. Of the 98 SP11 sequences, the type with eight cysteine residues was the most prevalent; thus, these eight residues were designated as disulfide bonds at positions 1-8, 2-5, 3-6, and 4-7. For *Br S*_46_-SP11 with seven Cys residues and *S*_32_-/*S*_36_-SP11 with ten Cys residues, the pairs were inferred from a previous study^*36*^. The other SP11s whose sequences closely resembled them were assumed to comprise the same pairs. After the application of CysBar, the 98 sequences were processed using Clustal Omega^*75, 76*^ to obtain “cys-seq” MSA. For comparison, “seq” MSA was similarly obtained without using CysBar. The resultant “seq” and “cys-seq” MSAs are shown in **Fig. S1**.

Additionally, two structure-based MSAs, namely, “st” and “st2” MSAs, were constructed using the 98 sequences for comparison with the sequence-based alignments. The St MSA was constructed as follows:

1. The two crystal structures for SP11 *S*_8_ and *S*_9_ and 16 modeled SP11 structures for *Br S*_12_, *S*_21_, *S*_25_, *S*_29_, *S*_32_, *S*_34_, *S*_36_, *S*_40_, *S*_44_, *S*_45_, *S*_46_, *S*_47_, *S*_49_, *S*_52_, *S*_60_, and *S*_61_, some of which were reported in our previous study^*36*^, were subjected to Mustang 3.2.3^*77*^ to obtain their structure-based MSA.
2. Each of the remaining 80 sequences were added to the MSA by creating pairwise alignments with one of the 18 sequences that they were most closely related to according to the phylogenetic tree (**Fig. S2**).

St2 MSA was created as follows:

1. First, all 98 SP11 structures were predicted using ColabFold with the “default” MSA.
2. Second, the structures were aligned using Mustang^*77*^.

Each of the four MSA files comprised 98 sequences. We ran the structure prediction using the MSA file with LocalColabFold by moving the corresponding sequence to the top of the file and reformatting it into the a3m format using the “reformat.pl” script, which is bundled in the HH-suite^*42*^. No templates were used for predictions using the four MSAs.

The normalized number of effective sequences (*N*_eff_) was computed for each position of a query sequence at a threshold of 80% sequence identity to estimate the quality of the MSA, which is also described as MSA depth in AlphaFold2^*38*^.

### Improvement of the prediction of class-II SP11

For class-II SP11s, a more confident model was predicted using ColabFold with a distinct MSA composed of class-II SP11 sequences retrieved from the BLASTP search as the input and a predicted structure that showed the best pLDDT among the four MSAs as the template. The recycling number for the prediction was increased to 9.

### Evaluation of predicted models

We used the pLDDT confidence measure^*38*^, which is a predicted per-residue lDDT-Cα score^*78*^, to evaluate the predicted monomeric model of eSRK or SP11. Additionally, we used the metrics—pTM-score^*38*^ and pDockQ^*79*^—to evaluate the predicted model for the eSRK–SP11 complex. The pLDDT and pTM-scores were calculated automatically using ColabFold for each predicted model, and we ranked the models according to the mean pLDDT for all residues per model. The pDockQ score was calculated using “pDockQ2.py” available at https://gitlab.com/ElofssonLab/huintaf2/-/blob/main/bin/pDockQ2.py and averaged for the chains in the model.

### Paired MSA for the prediction of eSRK–SP11 complex

To run the structure prediction for a complex with proteins A and B, ColabFold requires an a3m-formatted MSA file in which a horizontally concatenated MSA of protein pairs A and B that bind correctly is placed in the first block, followed by the MSAs of proteins A and B. In this study, we prepared an MSA for each of the 98 Clustal Omega-created eSRK sequences^*75, 76*^ and for the four types of SP11 sequences described in the previous section. We concatenated the aligned eSRK and SP11 sequences for each *S* haplotype in the “paired” block of the a3m file. Aligned homologous sequences of eSRK and SP11 with gaps were placed in the second and third blocks, respectively. The format of the input a3m file is shown in **Fig. S3**.

### MM–GBSA analysis

Molecular mechanics with generalized Born and surface area solvation (MM–GBSA)^*80*^ implemented in AmberTools 22^*81*^ was used to calculate the binding free energy, *ΔG*_bind_, between the modeled eSRK–SP11 complexes. The initial coordinates were obtained from a predicted complex model that exhibited the best pDockQ values for each haplotype. The ff19SB force field^*82*^ was used for the proteins, and the system was solvated using the OPC water model^*83*^. Energy minimization, equilibration, and a 50-ns conventional production run were performed using AMBER 22^*81*^ as per the procedure described in our previous paper^*36*^. The GBOBC implicit solvent model (parameters α = 1.0, β = 0.8, and γ = 4.85)^*84*^ was used with a salt concentration of 0.2 M. All calculations were performed using MD trajectories between 10 and 50 ns and recorded every 100 ps (400 snapshots for each complex).

## RESULTS

### eSRK and SP11 structures predicted using “default” MSA

We first used the MSAs retrieved from ColabFold/MMSeqs2 webserver, denoted as the “default” MSA, to predict the structures of eSRK and SP11 (**Figure 1A**). Of the 98 haplotypes, 97 predicted eSRK structures and 53 predicted SP11 structures exhibited an average pLDDT of ≥ 75. The high pLDDT of eSRKs could be attributed to their long amino acid sequence length (approximately 400) and high sequence homology between the haplotypes, whereas the low pLDDT of SP11 could have been caused by the short amino acid length and many diverse insertions and deletions in the defensin-like SP11 domain in the MSA, making the prediction difficult. The relatively low pLDDT observed in *Rs S*_22_-, *Rs S*_26_-, and *Rs S*_30_-SRK were due to the incomplete N- or C-terminal sequences (**Fig. S4**). **Figure 1B** shows the correlation between *N*_eff_ and the average pLDDT of SP11 structures predicted using the default MSA. The average pLDDT increased as *N*_eff_ approached 100, which is consistent with the discussion of AlphaFold2^*38*^. However, five outlier proteins, namely, *Br S*_12_-, *Br S*_33_-, *Rs S*_23_-, *Rs S*_31_-, and *Bo S*_11_-SP11, exhibited low pLDDT despite having *N*_eff_ > 500 because most of the MSAs are composed of inappropriate amino acid sequences that are partially analogous but not homologous to the full-length SP11. In contrast, only a limited number of homologous sequences could be obtained from the webserver for *Br S*_36_- and *Bo S*_24_-SP11, which was insufficient to obtain a successful prediction. The left panels of **Fig. S5A** and **B** visualize the MSA coverage of the default MSA for *Br S*_12_- and *Br S*_36_-SP11, respectively.

**Figure 1.**
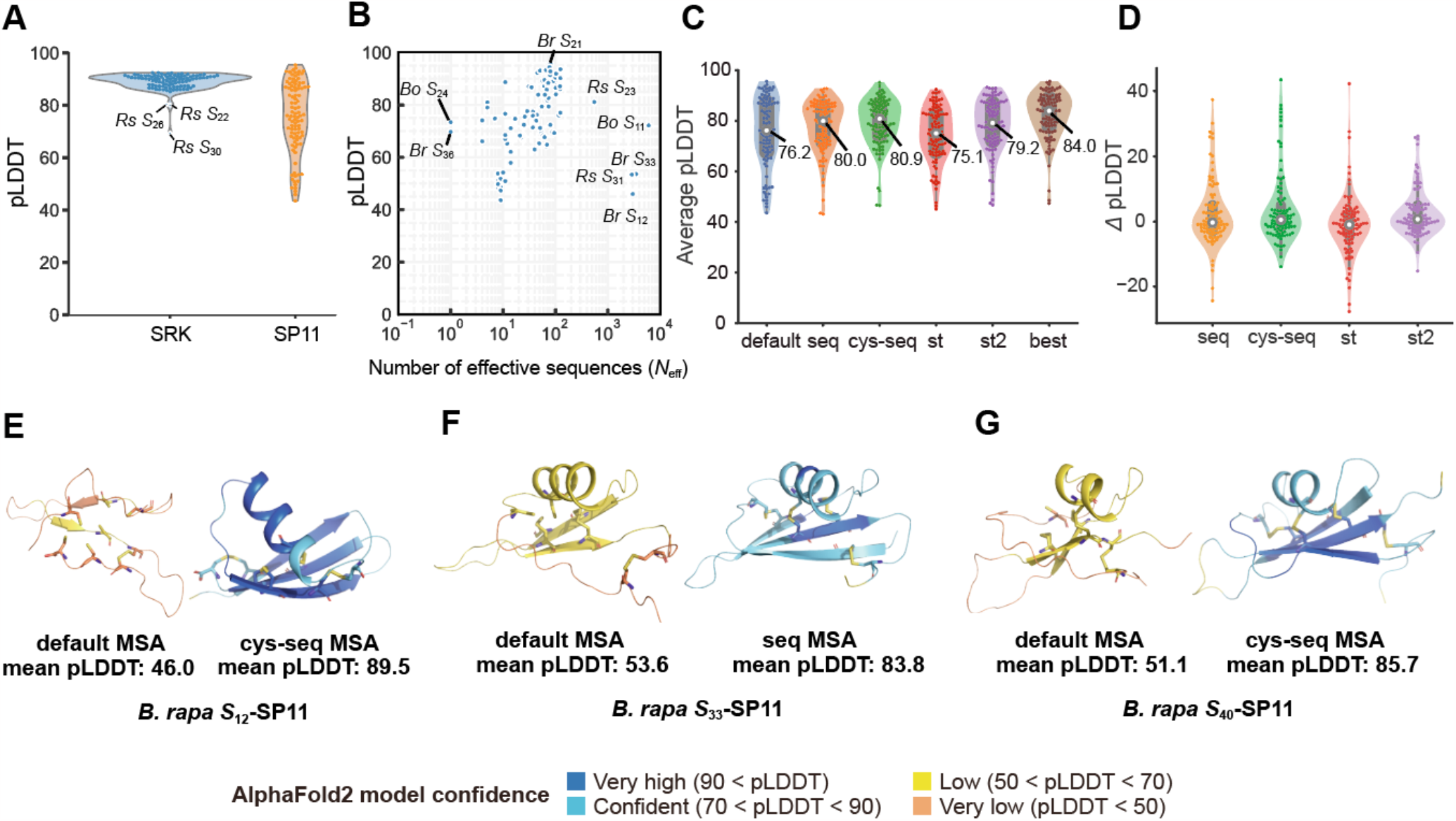
Structure prediction of S-locus protein 11 (SP11) proteins with ColabFold using various multiple sequence alignments (MSAs). (A) The mean predicted local distance difference test (pLDDT) of *S* receptor kinase (SRK) and SP11 models predicted using ColabFold with the “default_MSA” (uniref30_2202 + bfd_mgy_colabfold database). (B) The mean pLDDT of SP11 models for the MSA depth. *N*_eff_ was calculated according to a previously reported procedure^*94*^ with a threshold of 62% sequence identity. (C) The mean pLDDT of the predicted SP11 models vs. the four custom MSAs (*n* = 98). The white circle represents the median. (D) Difference in pLDDT with respect to the default MSA. The white circle represents the median. (E–G) Significantly improved models of *Br S*_12_ (E), *Br S*_33_ (F), and *Br S*_40_ (G). Models predicted with the default and one of the four MSAs which exhibits the best pLDDT are shown in left and right panels, respectively.

### Custom MSA improved SP11 models

ColabFold enables other MSA files to be specified as input instead of the default MSA. To obtain improved SP11 model structures, we prepared four types of MSA and denoted them as “cys-seq”, “seq”, “st”, and “st2”; each consisted of 98 SP11 sequences that were experimentally annotated (see Methods). All four custom MSAs yielded SP11 models with a pLDDT score equivalent to or higher than that of the default MSA (**Figure 1C and D**). For 66 of the 98 SP11s, the prediction accuracy was improved when one of the four MSAs was used. The median pLDDT increased to 84.0 by choosing the best of the four MSA models, and the number of SP11s showing pLDDT > 75 increased from 53 to 81. Particularly, the *Br S*_12_-SP11 model, which was predicted using cys-seq MSA, showed the largest improvement (46.0 to 89.5 in the mean pLDDT; **Figure 1E**). Similarly, the prediction with custom MSAs also improved for *Br S*_33_, *Bo S*_11_, *Rs S*_23_, and *Rs S*_31_ (**Fig. S6**).

Although the four MSAs improved the pLDDT scores for many SP11s, they had little effect for *Rs S*_11_ (48.5), *S*_15_ (52.3), and *S*_26_ (47.3) because their C-terminal sequences after the fifth Cys residue corresponding to the β3 strand are missing (**Fig. S1**). Similarly, the pLDDT scores of *Br S*_46_, *S*_61_, *Bo S*_7_, *S*_13_, and *S*_13b_ were < 70 despite their sequences being close to *Br S*_8_, the crystal structure of which was determined. All pLDDT scores for the predicted models obtained using the default and four MSAs are provided in **Supplementary File 2**.

### Structural novelty in the predicted SP11 proteins

Our predicted models with high pLDDT values enable us to explore the polymorphisms of SP11 proteins based on their sequences and structures. Although most SP11s typically possess eight Cys residues and the disulfide pairs are conserved to be 1-8, 2-5, 3-6, and 4-7^*85*^, we found three structural groups with different disulfide bond pairs in SP11s with 10 Cys residues. First, *Br S*_32_ and *Br S*_36_ showed 1-10, 2-6, 3-7, 4-9, and 5-8 disulfide bond connectivity (**Figure 2A–B**), in which two disulfide bonds exist between the β2 and β3 strands. The bond connectivity and regions of the secondary structures of *Br S*_32_ and *Br S*_36_ were accurately predicted in our previous study using homology modeling and accelerated MD simulations^*36*^ (**Fig. S7**). The predicted SP11 models of *Bo S*_24_ and *Bo S*_68_ also exhibited the same connectivity, and they closely resembled *Br S*_36_ and *Br S*_32_, respectively, because their sequence similarities were substantially high (**Fig. S2**). Second, *Br S*_54_, *Bo S*_28_, and *Rs S*_31_ exhibited 1-9, 2-10, 3-6, 4-7, and 5-8 disulfide bond connectivity, in which two Cys residues on the β1 strand form bonds with those located after the β3 strand (**Figure 2C–E**). Notably, SP11 of *Rs S*_31_ formed a β4 strand at its C-terminus (**Figure 2E**). Third, *Br S*_26_-SP11 showed the 1-10, 2-5, 3-7, 4-8, and 6-9 connectivity (**Figure 2F**). It possesses two helices between the β1 and β2 strands, although the second helix is not involved in the interaction with its corresponding eSRK (**Fig. S8**).

**Figure 2.**
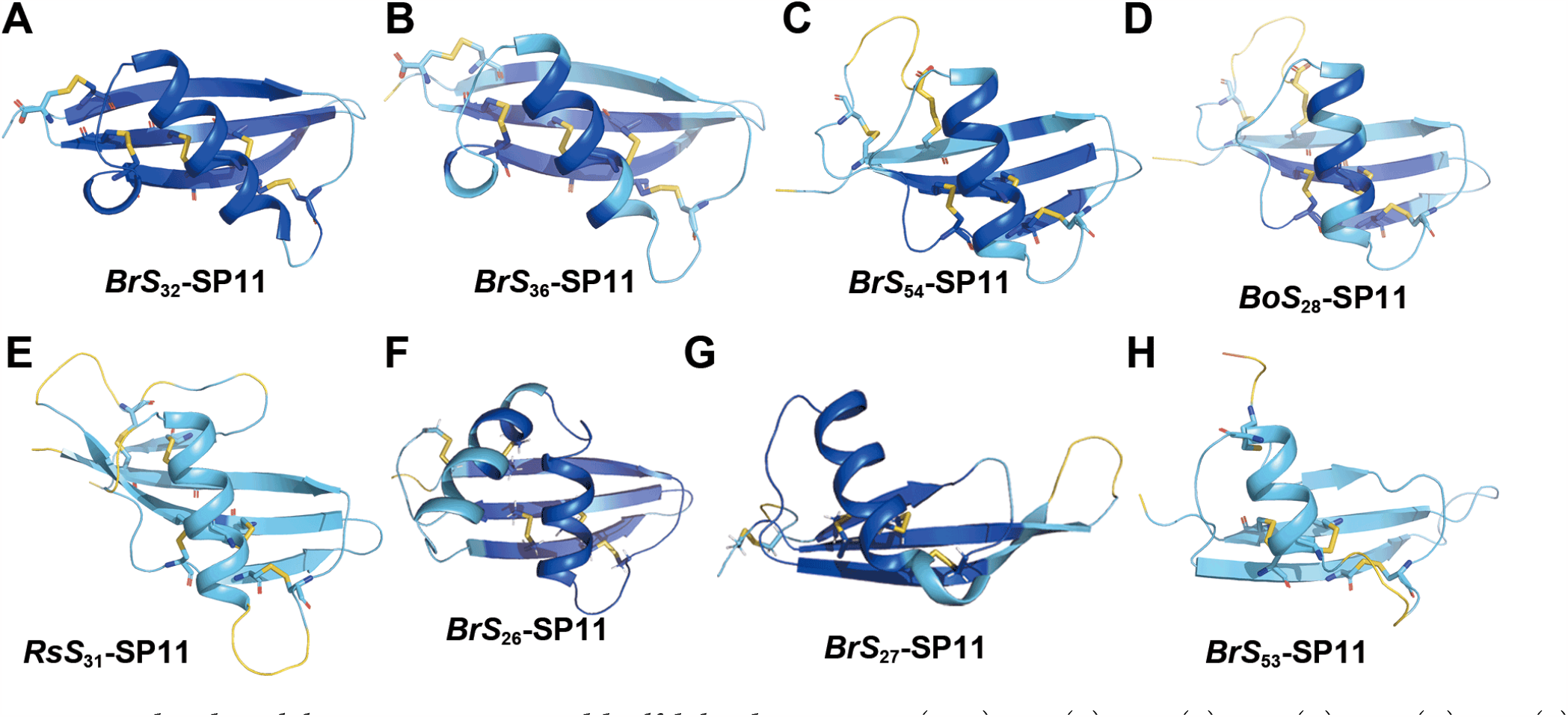
Predicted novel class I SP11 structures and disulfide bond connectivity. (A–H) *Br S*_32_ (A), *Br S*_36_ (B), *Br S*_54_ (C), *Bo S*_28_ (D), *Rs S*_31_ (E), *Br S* (F), *Br S*_27_ (G), *Br S*_53_ (H)-SP11 model structures predicted using ColabFold with custom MSAs. Disulfide bonds are shown as sticks. The models are colored according to the AlphaFold2 model confidence.

Furthermore, two anomalous structures are worth mentioning in the SP11s with eight Cys residues: the first is *Br S*_27_ and its one-residue variant, *Bo S*_8_; both exhibited long twisted β-hairpin structures (**Figure 2G**). The second is *Br S*_53_, which exhibited a structure comprising 1-4, 2-6, 3-7, and 5-8 disulfide connectivity (**Figure 2H**); the fourth Cys residue was located at the end of the α helix.

Of the class-II 10 *S* haplotypes, *Br S*_29_, *Br S*_40_, *Br S*_44_, *Br S*_60_, *Bo S*_5_, *Rr S*_6_, and *Rs S*_9_ exhibited a mean per-residue pLDDT of ≥ 75 (**Figure 3A–G, middle**), and their backbone structures resembled each other (**Figure 3H**). The prediction for the remaining three haplotypes, *Rs S*_11_, *Rs S*_15_, and *Rs S*_26_, still failed for the reason described in the previous section. Applying our refinement protocol for class-II SP11s (see Methods) to the seven haplotypes with accurately predicted structures improved their pLDDT values to > 87 (**Figure 3A–G, right**). The predicted class-II structures possessed a characteristic β-bridge secondary structure after the first β-strand (**Figure 3I and J**). The α-helix of *Br S*_60_ was longer than that of these haplotypes by two residues (**Figure 3D**). Remarkably, only the *Rs S*_9_ model lacked the α-helix between the β1 and β2 strands among the 98 SP11s (**Figure 3G**) regardless of its high mean pLDDT score (91.7). All seven predicted structures showed substantially higher pLDDT values than of those presented in the AlphaFold DB version 2022-11-01 (**Figure 3A–G, left**), indicating that new predicted structures were more plausible. The low confidence of the models in the AlphaFold DB could be attributed to the failure to detect the class-I SP11 sequences, which are evolutionarily distant from those of class-II sequences and to the inappropriate alignment of cysteine-rich protein fragments during the construction of the MSA.

**Figure 3.**
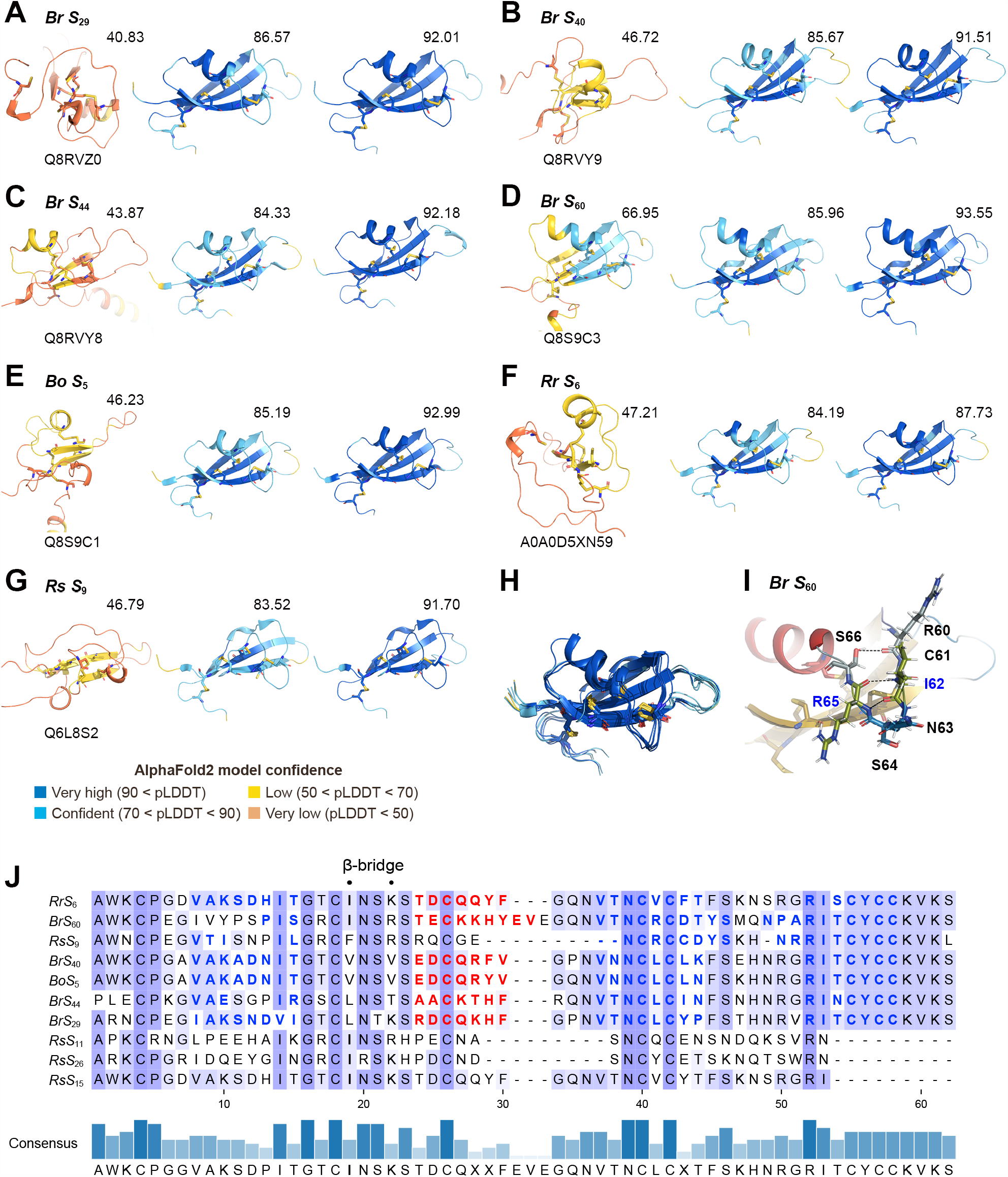
Predicted models of class-II SP11s. (A–G) Predicted SP11 models of *Br S*_29_ (A), *Br S*_40_ (B), *Br S*_44_ (C), *Br S*_60_ (D), *Bo S*_5_ (E), *Rr S*_6_ (F), and *Rs S*_9_ (G) haplotypes. The model deposited in AlphaFold DB (version 2022-11-01), the best one among the four MSAs, and one improved using the refinement protocol are depicted in the left, middle, and right panels, respectively. UniProt accession IDs are shown under the model of AlphaFold DB. The structures are colored according to the AlphaFold pLDDT model confidence. The mean per-residue pLDDT is shown above each model on the upper right side. (H) Superposition of the seven predicted class-II SP11 proteins. (I) A characteristic β-bridge structure observed in the predicted class II SP11 proteins. *Br S*_60_-SP11 is shown as an example. The hydrogen bonds are shown as yellow dashed lines. (J) Sequence alignments of class-II SP11 proteins. The α-helix and β-strand secondary structures annotated using DSSP 4.0.5 are shown as red and blue characters, respectively. The characteristic β-bridge are indicated using “·” marks.

### Improved eSRK–SP11 complex prediction using ColabFold and paired MSAs

We also investigated the effect of the four MSAs on the modeling of the eSRK–SP11 complexes. We used three criteria to assess the plausibility of the predicted eSRK–SP11 complex models: the pTM-score^*38*^, pDockQ score^*79*^, and a root-mean-square-deviation (rmsd) for the crystal structure of the *Br S*_8_ eSRK–SP11 complex (PDB:6KYW). Of the 98 predicted complex models with their default MSAs, only 41 showed pDockQ > 0.6. However, the prediction with a paired MSA, where one of the four SP11-MSAs are concatenated with the eSRK-MSA (see **Fig. S3**), led to 80 models meeting the criteria. (**Figure 4A**). Notably, st2 and seq MSAs significantly enhanced the pDockQ scores of *Br S*_34_ and *S*_55_ complexes, respectively. Consequently, their two SP11 molecules were located between the eSRK dimers, similar to the conformation observed in the *Br S*_8_/*S*_9_ crystal structure (**Figure 4C and D**). Similarly, *Br S*_12_ and *Br S*_47_ complex models were also improved by cys-seq and st MSAs, respectively (**Figure 4E and F**). Overall, the four paired MSAs led to an enhanced pDockQ score for 95 complex models (**Table 1**), and the most effective MSAs varied depending on the *S* haplotype (see **Supplementary File 2**).

**Table 1.** Evaluation of each paired MSA used for predicting the eSRK–SP11 complex.

|  | mean pDockQ<sup>a</sup> | mean rmsd [Å]<sup>a</sup> | Num. of best pDockQ structure |
| --- | --- | --- | --- |
| default | 0.424 | 5.06 | 3 |
| seq | 0.623 | 3.39 | 24 |
| cys-seq | 0.616 | 3.75 | 24 |
| st | 0.591 | 3.59 | 15 |
| st2 | 0.600 | 3.96 | 32 |

<sup>a</sup> average of the 98 predicted models.

**Figure 4.**
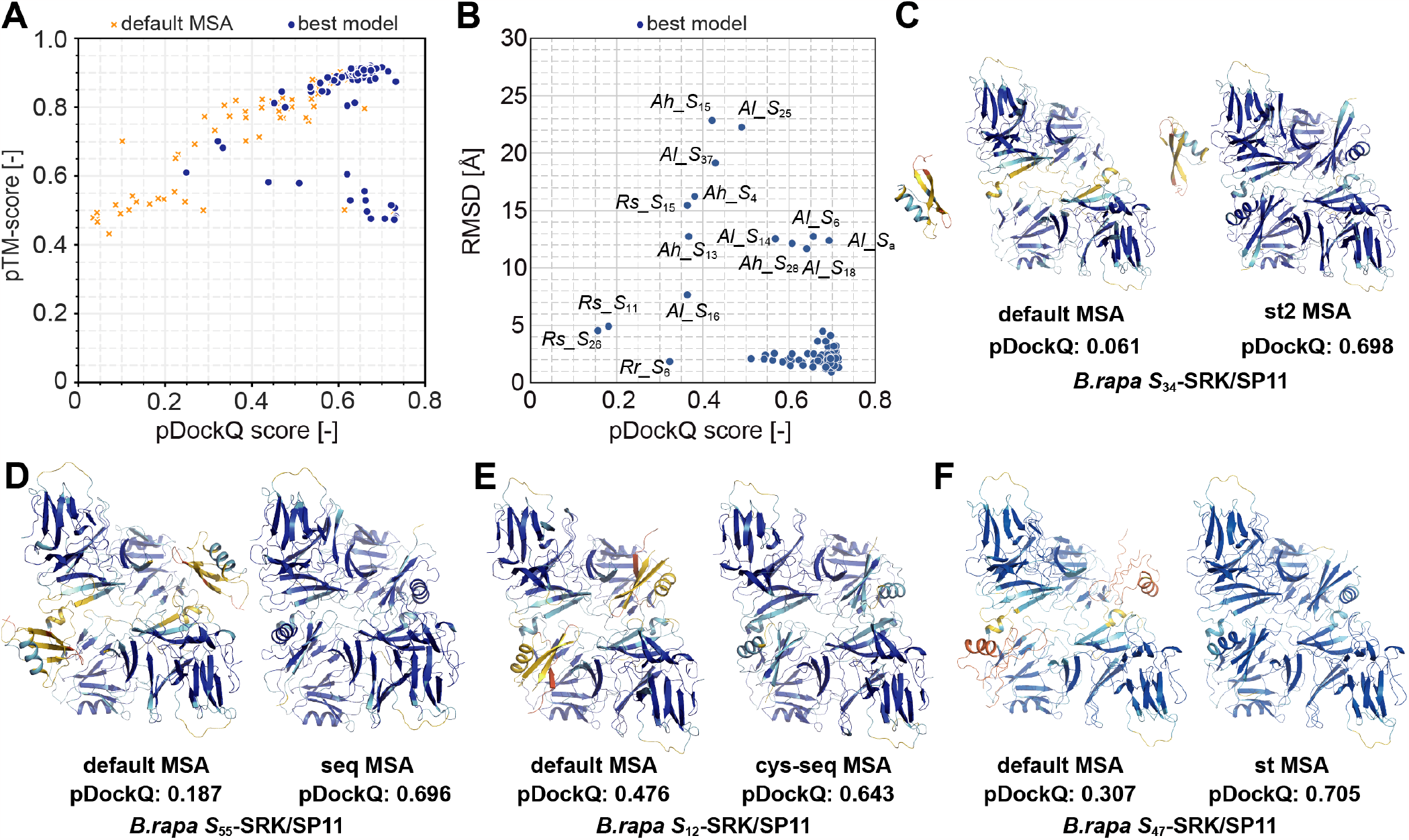
Complex predictions with the custom MSAs. (A) Correlation between pTM-score and pDockQ for the predicted complex models. Orange crosses and blue circles indicate the complex models predicted using the default MSA and the highest pDockQ of the four MSAs, respectively. (B) Root mean square deviation (RMSD) values of the predicted model showing the highest pDockQ (best model) to the *Br S*_8_ crystal structure (PDB: 6KYW). (C–F) Improved complex models of *Br S*_34_ (C), *Br S*_55_ (D), *Br S*_12_ (E), and *Br S*47 (F) haplotypes using the custom MSAs. The residues are colored based on the pLDDT values.

Of the 80 improved complex models, 76 exhibited rmsd < 5 Å, indicating that they share the same binding mode as that of *Br S*_8_ or *Br S*_9_. The complex predictions for *Rs S*_11_, *Rs S*_15_, *Rs S*_26_ were not successful owing to their poor SP11 models as described earlier. Remarkably, among the prediction for *Ah* and *Al, Ah S*_28_, *Al S*_6_, *Al S*_14_, *Al S*_18_, and *Al S*_a_ (also known as *Al S*_13_) showed rmsd > 10 Å despite pDockQ > 0.50, although their monomeric structures were predicted with high pLDDT values (**Figure 4B**). They formed a novel binding interface in which an SP11 molecule was placed between HV-II of one eSRK and HV-III of the other eSRK (**Figure 5A**). In contrast, *Ah S*_12_, *Ah S*_20_, *Ah S*_32_, *Al S*_36_, and *Al S*_b_ (also known as *Al S*_20_) showed a complex mode similar to that of the *Br S*_8_ or *Br S*_9_ crystal structures, in which HV-II was in direct contact with HV-III (**Figure 5B**). The predicted aligned error (PAE) indicates the unreliability of the residue–residue relationship of the predicted AlphaFold2 model^*38*^. PAE was generally low for both models (**Figure 5C and D**), suggesting that these predicted models were both plausible.

**Figure 5.**
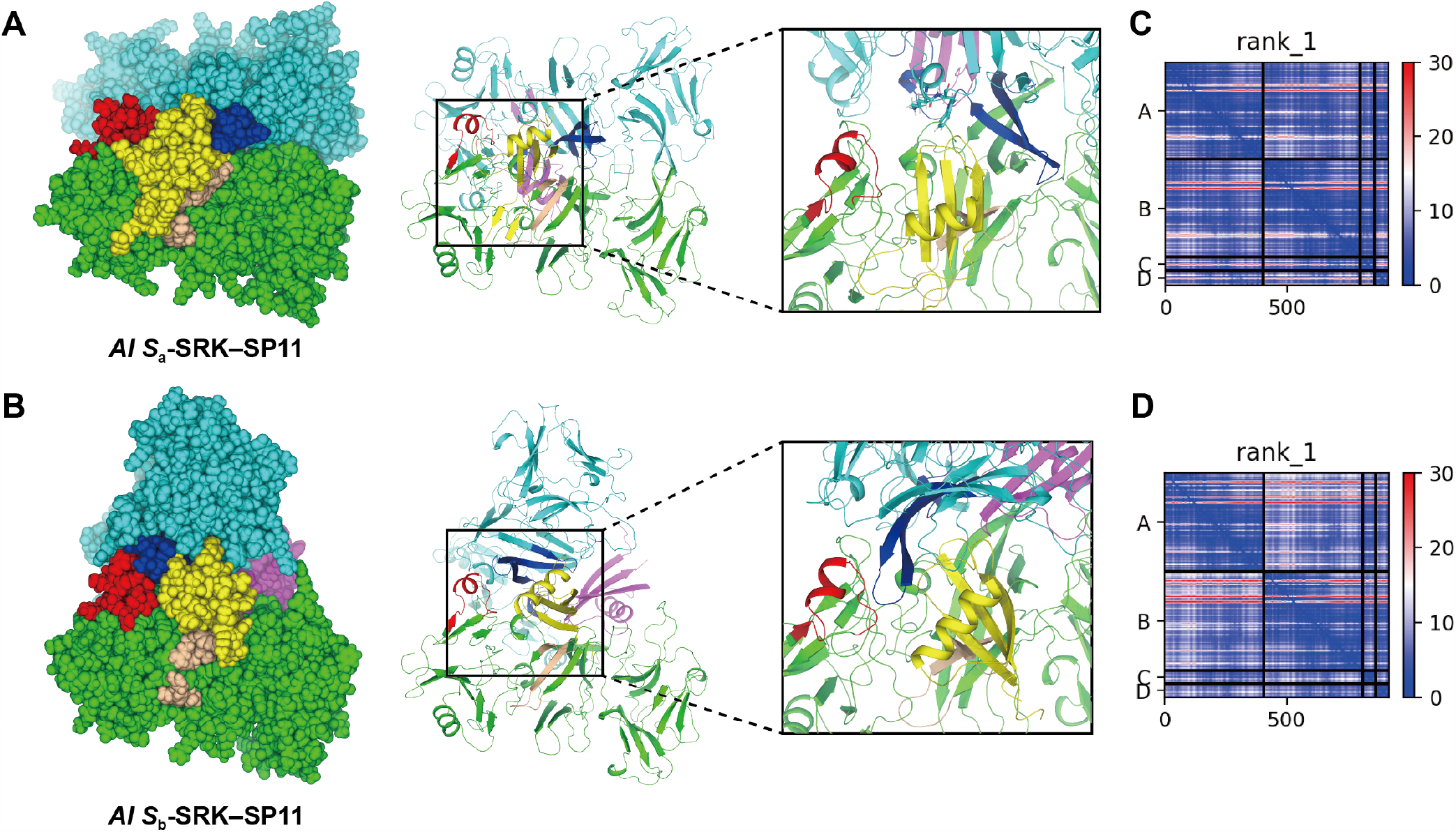
redicted binding mode for some *Al* or *Ah* SRK–SP11 complexes. (A–B) Predicted *Al S*_a_ (A) and *Al S*_b_ (B) complex models using ColabFold. Two SRK monomers are colored in green and cyan, and two SP11s are in yellow and purple. The position and orientation of *Al S*_a_- and *Al S*_b_-SRK chain A (green) are aligned. The HV regions I, II, and III of SRK are colored in light brown, deep blue, and red, respectively. HV-I and III belong to the same SRK chain, whereas HV-II belongs to the other chain. Sphere and cartoon representations are shown in the left and center panels, respectively. A close-up view of the binding interface between SRK and SP11 are shown in the right panel. (C–D) The predicted aligned error (PAE) for *Al S*_a_ (C) and *Al S*_b_ (D). Chains A–B and C–D represent SRK and SP11, respectively.

### Binding free energy calculations for the predicted eSRK– SP11 complexes using MM–GBSA analysis

In this study, we obtained highly plausible eSRK–SP11 complex models for most haplotypes using ColabFold and curated MSAs. However, they do not guarantee tight interactions because the AlphaFold2/ColabFold prediction is statistical and not physical. Hence, we used MM–GBSA analysis^*80*^ to calculate the binding free energy (*ΔG*_bind_) for the predicted models.

Of the 98 complex models that were examined, 57 structures demonstrated *ΔG*_bind_ < –75 kcal/mol. **Figure 6A** illustrates the per-residue contribution to *ΔG*_bind_ on a sequence alignment of class-I *Br* SP11s arranged using CysBar^*72*^ and Clustal Omega^*75, 76*^. In general, the residues comprising the first loop region, the α helix, and the β2-β3 hairpin exhibited large negative *ΔG*_bind_, which conforms with the results of our previous study^*36*^ and the binding interfaces observed in the predicted complex models. The MM–GBSA analysis showed largely negative *ΔG*_bind_ values for haplotypes whose SP11 showed a characteristic secondary structure; in *Br S*_26_, the contribution of *ΔG*_bind_ to eSRK interactions was observed near the kink between the first and second helices (**Fig. S8**); in *Br S*_27_, another SP11 molecule interacts with its unique β2–β3 structure. These results demonstrate that most of the predicted complex models are truly bound with high affinity. The per-residue energy contributions for all haplotypes that showed a large negative *ΔG*_bind_ in this study are shown in **Supplementary File 3**.

**Figure 6.**
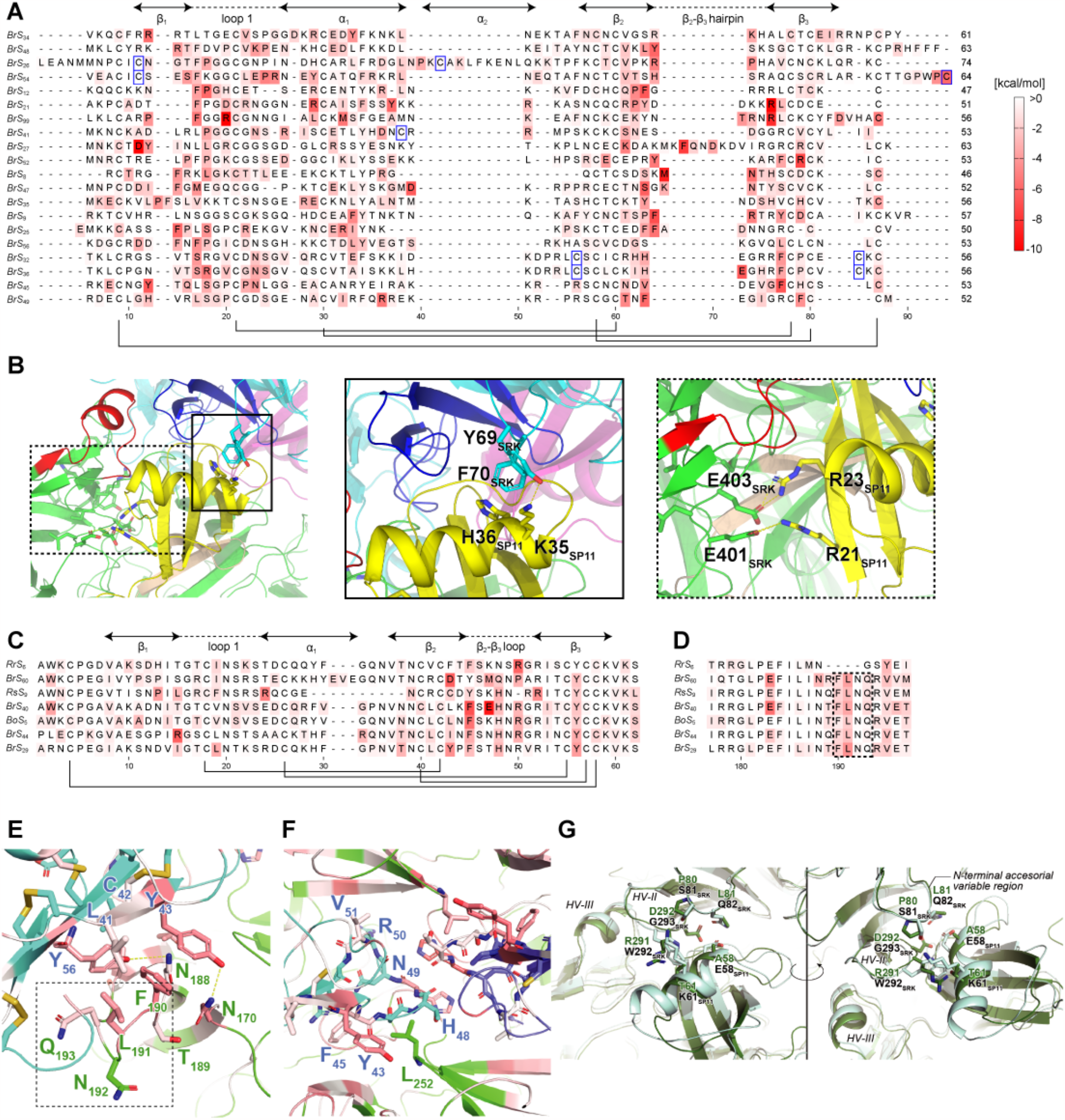
Molecular mechanics–generalized born surface area (MM–GBSA) analysis using the predicted structures of SRK–SP11 complexes. (A) *Br* class-I SP11 sequences with heat maps indicating energy contributions from each residue. Cys residue pairs forming disulfide bonds are shown in black lines. Extra Cys residues observed in class-I SP11s are highlighted in blue squares. (B) N- and C-terminal accessorial variable regions of eSRK in the *Br S*_48_ SRK–SP11 complex model. Close-up views of the N- and C-terminal regions are depicted in the middle (solid line) and right (dashed line) panels, respectively. The color scheme of the protein model is the same as in **Fig. 1**. (C) The heat map of *Br* class-II SP11 sequences. (D) The heatmaps of HV-I of class-II ectodomain SRKs (eSRKs). The FLNQ insertion are indicated by the dashed lines. (E–F) Close-up view of the binding interface around the eSRK HV-I region (E) and the SP11 β2-β3 loop (F) in the predicted *Br S*_29_ model. eSRK and SP11 are colored as cyan and green, respectively. Residues at the interface are colored according to their per-residue *ΔG*_bind_ values. Subscripts next to the one-letter residue name indicate the residue number in the sequence alignment. (G) Superposition of predicted *Br S*_44_ (green) and *S*_60_ SRK–SP11 (pale blue) complex models.

Based on the predicted complex structures and MM–GBSA analysis, we propose the presence of two additional variable regions in eSRK: the N- and C-terminal accessorial variable regions. These regions are partly involved in the self/nonself-discrimination mechanism. For example, in *Br S*_48_, Tyr69 and Phe70 that are located in the N-terminal accessorial variable region interacted with the α-helix of SP11, and Glu401 and Glu403 that are located in the C-terminal region formed salt bridges with Arg21 and Arg23 of SP11, respectively (**Figure 6B**). Importantly, these regions do not necessarily contribute to the eSRK–SP11 interactions depending on the haplotype, as was not observed in the crystal structures of *Br S*_8_ and *S*_9_ (see also **Supplementary File 3**). Particularly, the C-terminal variable region was previously discovered by Kusaba *et al*. but it was not believed to be involved in the specificity between SRK and SP11^*29*^. However, the predicted models and analyses suggest that this contributes to complex stabilization in some *S* haplotypes.

The predicted class-II complex structures presented a different binding mode from that observed for class-I. The amino acid compositions of β3 in class-II SP11s are well conserved, and a conserved Tyr residue at position 56 of SP11 interacted with the insertion characteristic of HV I in class-II eSRK, except in *Rr S*_6_ (**Figure 6C–E and Figure 3H**). Notably, the loop between β2–β3 of class-II SP11s interacted with that of another SP11 molecule in the class-II complex models (**Figure 6F**). Moreover, unlike that observed in the class-I SP11 models such as *Br S*_8_ and *Br S*_9_, the α-helix in class-II did not interact with HV III of the corresponding eSRKs (**Fig. S9**), indicating that the HV III of eSRK of class-II does not contribute to the *ΔG*_bind_.

The high similarity of the backbone structure of the predicted SP11 and eSRK–SP11 complex models in the class-II models, except for *Rr S*_6_, and the well-conserved amino acids consisting of the β3 strand of SP11 suggest that the self/nonself-discrimination within the class II is achieved by the other binding interfaces. Structural comparison of the predicted eSRK–SP11 complexes of *Br S*_44_ and *Br S*_60_ suggested that the slightly bulky Glu58 and Lys61 of *S*_60_-SP11 are located in a manner so as not to interfere with Trp292 and Gly293 of *S*_*60*_-eSRK, whereas in the *S*_44_ model, Ala58 and Thr61 of SP11 and bulky Arg291 and Asp292 of eSRK are located at the same position (**Figure 6G**). According to the sequence alignment of the class-II eSRKs and SP11s (**Figure 3H**), only *S*_44_ exhibited an inversion of bulkiness, suggesting that it plays a key role in the self/nonself-discrimination in class II and in the N-terminal accessorial variable region.

Interestingly, the eSRK of *Rr S*_6_ lacks the four-residue FLNQ insertion characteristic of the class-II HV-I region, whereas its SP11 displayed a class-II-like sequence and predicted structure (**Figure 6D**). The pDockQ score of the predicted *Rr S*_6_ complex model was not highly plausible (pDockQ = 0.322) compared with that of the other class-II predicted models (mean pDockQ = 0.588). Hence, we independently superimposed the predicted *Rr S*_6_ eSRK and SP11 models onto the predicted *Br S*_29_ complex structure to generate a new complex model equivalent to the other class-II complex models showing high pDockQ. Using the model as the initial coordinate of MD simulations, the MM–GBSA analysis demonstrated that the calculated Δ*G*_bind_ between the eSRK and SP11 models was negative (–36.3 kcal/mol) and identified the residues contributing to the binding. Analyzing *Rr S*_6_ showed that the Tyr residue at position 56 of SP11 did not contribute to the Δ*G*_bind_ (**Figure 6C**). Taken together, these observations suggest that binding and self-recognition in class II haplotypes are predominantly mediated by the interaction between the α-helix of SP11 and the HV-II/N-terminal accessorial variable region of eSRK. Additionally, the interaction between the Tyr residue at position 56 of SP11 and the FLNQ insertion of eSRK augments the affinity.

Finally, we also evaluated the five haplotypes showing the different binding mode using MD simulations. The *Ah S*_28_, *Al S*_6_, *Al S*_14_, and *Al S*_a_ complex models exhibited *ΔG*_bind_ values of –139, –88.3, –60.3, and –98.7 kcal/mol, respectively, indicating that they were plausibly modeled and their cognate eSRK and SP11 molecules were tightly bound. In contrast, *Al S*_18_ showed *ΔG*_bind_ = –12.6 kcal/mol, suggesting that the model was partially inaccurate. Next, we counted the number of atomic contacts between the eSRK and SP11 molecules in the modeled *Ah S*_28_ and compared it with that of the model based on the *Br S*_9_ crystal structure geometry^*86*^. The *Ah S*_28_ SRK-SP11 complex model showed a lower number of atomic contacts (1251 contacts) than that of the previously modeled one (1395 contacts) but higher than those of the *Br S*_8_ or *Br S*_9_ crystal structures (1180 and 1159 contacts, respectively, **Table S1**). Moreover, the Arg277 and Glu279 of chain A in the *Ah S*_28_-eSRK model formed salt bridges with those in chain B (**Figure 7A**). Additionally, Arg218 of the eSRK model formed an H-bond network with Asp288/Glu293 of the other chain of the eSRK and Arg68 of the cognate SP11 (**Figure 7B**). Although these interactions were not observed in the previous model, our contact number analysis and *ΔG*_bind_ calculation suggest that the different binding mode is also probable.

**Figure 7.**
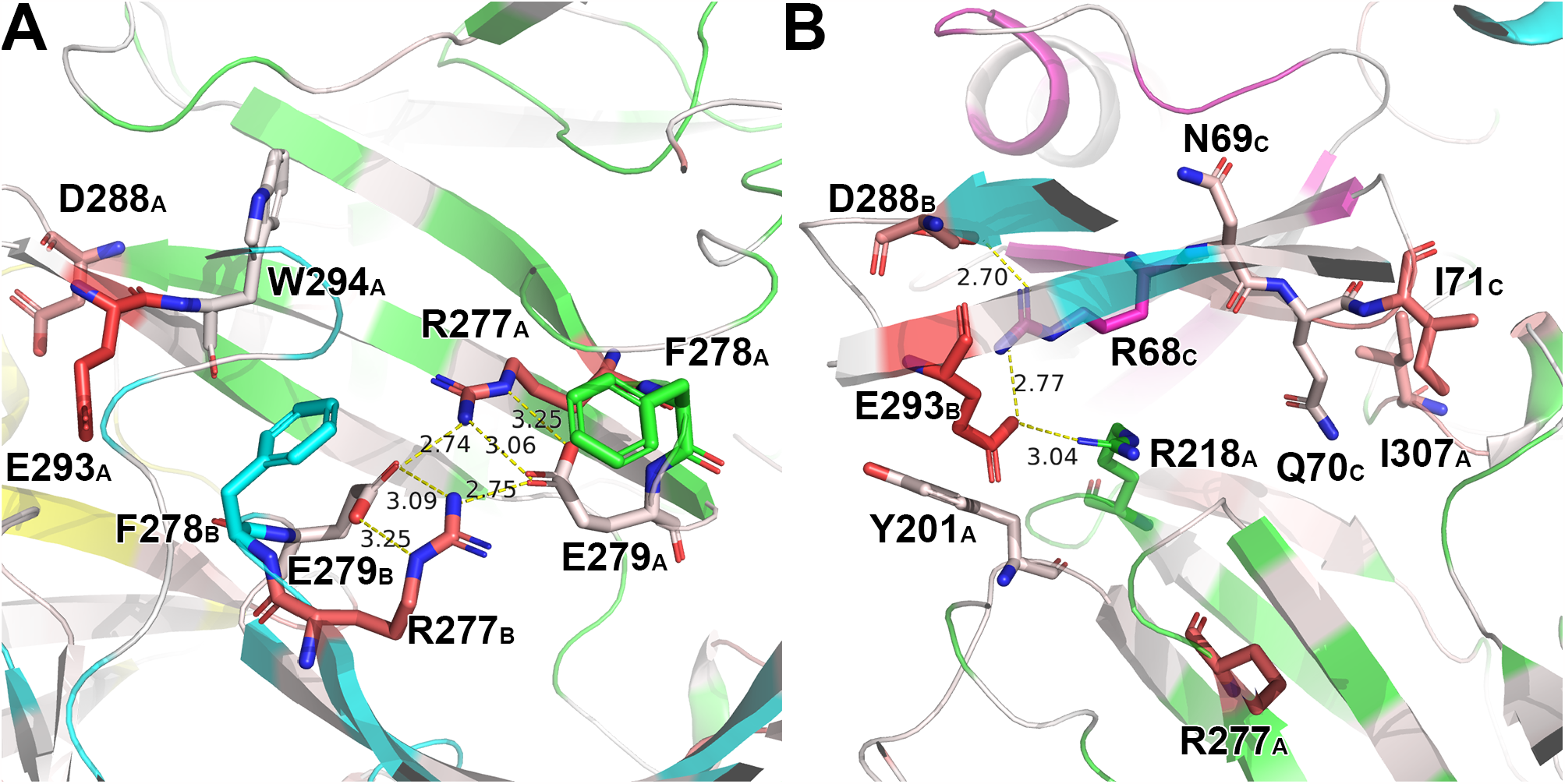
The binding interface of *Ah S*_28_ eSRK–SP11 complex model. Chain A (green) and B (cyan) belong to eSRK and Chain C (purple) and D (yellow) to SP11. The dashed line represents hydrogen bonds. Bond lengths are shown in angstroms. Residues are colored according to their per-residue ΔGbind values as in Fig. 6. The residue numbers are labeled in conformance with the GenBank database. (A) The interface between the two eSRK molecules. (B) eSRK–SP11 interface.

## DISCUSSION

In this study, we demonstrated that SP11 proteins from various species could be predicted with high pLDDT using ColabFold with manually curated and custom MSAs. We have also presented eSRK– SP11 complex models using paired eSRK and SP11 sequences originating from the same *S*-haplotype. The difficulty encountered in predicting the SP11 defensin-like domain structure using the original AlphaFold2/ColabFold framework stems from alignment errors pertaining to the presence of eight or more cysteine residues within its approximately 60-residue sequence. Here, we circumvented this issue by employing four custom MSAs grounded in the assumption that Cys residues form disulfide bonds with fixed connectivity. Lastly, a comprehensive analysis of the binding mechanism between eSRK and SP11s led to the discovery of novel specificity-determining regions of SRK located at the N- and C-terminal accessorial variable regions. We expect that the computational results will provide significant clues to a comprehensive understanding of the self/nonself-discrimination mechanism in Brassicaceae, which has been deemed challenging because of the polymorphism in sequence and structure.

The accuracy of AlphaFold2 relies on the extraction of coevolutionary information from the input primary sequence and its corresponding MSA generated in the calculation pipeline.^*38*^ The Evoformer block of AlphaFold2, inspired by direct coupling analysis (DCA)^*87-91*^, is tasked with predicting the inter-residue distances in the input sequence by iteratively refining the spatial coordinates and evolutionary information within the MSA. Importantly, the accuracy significantly drops if the median per-residue *N*_eff_ is less than approximately 30 but stabilizes when *N*_eff_ exceeds 100. Consequently, the quantity and quality of MSAs play an important role in the accuracy of the model. However, obtaining an accurate sequence alignment is difficult for small cysteine-rich proteins with few homologs in the sequence database. Our predictions for SP11s using ColabFold and the various MSAs (**Figure 1B**) also confirm the significance of *N*_eff_. Additionally, the improved model accuracy could be attributed to the rigorous preparation of the MSA, which comprises 98 pairs of coevolving eSRK and SP11 amino acid sequences along with the meticulously corrected cysteine residue positions. Overall, the eSRK–SP11 complex prediction for *Brassica* and *Raphanus* using our custom MSAs showed high pLDDT and pDockQ values, whereas those for some *S* haplotypes of *A. halleri* and *A. lyrata* were low. This result may be attributed to the small number of sequences that are closely related to the genus *Arabidopsis* in the sequence dataset.

Our MM–GBSA analysis revealed that the calculated *ΔG*bind values were largely negative for 57 cognate eSRK-SP11 pairs, indicating that their predicted models were physically plausible. We performed the analysis for cognate pairs in this study because the number of non-cognate pairs is substantially large. However, our models provide a reasonable explanation for interspecific pairs exhibiting the same recognition specificities such as in *Bo S*_7_–*Br S*_46_, *Bo S*_32_–*Br S*_8_, *Bo S*_24_–*Br S*_36_, *Bo S*_12_–*Br S*_47_, and *Bo S*_64_–*Br S*_41_ pairs^*68, 92*^. Their SRK and SP11 sequences have a > 92% sequence identity for each pair and superposition of their models showed that the mutated residues in each pair were not located around the eSRK–SP11 binding interfaces (**Fig. S10**).

Apart from the studies investigating the crystal structures of *Br S*_8_ or *S*_9_, few in vitro experimental studies in the past two decades have directly investigated the eSRK–SP11 molecular interactions because of the difficulty in expressing these proteins under laboratory conditions. Instead, the interaction has been analyzed through an in vivo assay using transgenic plants with mutation in their HV I-III regions. Boggs *et al*. constructed an eSRK chimera: eSRKa(7)a. Its *Al S*_a_ HV I-III regions were replaced with those of *Capsella grandiflora* (*Cg*) *S*_7_, which is 77% similar to *Al S*_a_. They demonstrated that the transgenic plant expressing eSRKa(7)a showed an incompatibility response to pollen expressing *Cg S*_7_-SP11. Additionally, they identified six single mutations of eSRKa(7)a that disrupt its function. We performed the prediction for *Cg* eSRK– SP11 to validate whether the results and our predicted models were consistent (**Fig. S11 and Supplementary File 4**). The predicted complex model exhibited almost the same binding mode as that of *Al S*_a_, and the six residues were located at the eSRK–SP11 interface. Moreover, the models also explained the adverse impact of each single mutation on the interaction. The K213M and Y301T mutations in eSRKa(7)a were predicted to disrupt the interactions formed by Thr66 and Asp69 in *Cg S*_7_-SP11, respectively, and the I217R mutant was predicted to interfere with Arg34 in *Cg S*_7_-SP11. Moreover, the backbone structure of residues 294–300, which is derived from *Cg S*_7_, was predicted to differ from *Al S*_a_ because of the exceptional phi torsion angle of Pro294. Thus, the P294S, X(gap)298A, and D300E mutations are predicted to alter the main chain structure and thereby change the incompatibility response. Meanwhile, a weakened response caused by the L218V mutation may be attributed to a slight decrease in binding affinity between eSRKa(7)a and *Cg S*_7_-SP11 owing to its smaller volume, although the model could not represent the effect well. Unfortunately, we could not examine the other chimera, eSRK16(25)16, in which they replaced the HV I-III regions of *Al S*_16_ with those from *Al S*_25_, due to failure in predicting their complex models.

Another crucial factor that needs to be mentioned is the relevance of eSRK–SP11 binding modes and the phylogenetic characterization for the allelic diversity of the *S*-locus in *Arabidopsis* and *Brassica* species. Prigoda *et al*.^*93*^ and Goubet *et al*.^*58*^ have suggested that *Ah S*_28_, *Al S*_6_, *Al S*_14_, and *Al S*_18_ belong to class B/II, and *Al S*_a_ (*S*_13_) is distinctly classified as class A3/III. Additionally, *Ah S*_12_, *Ah S*_20_, *Ah S*_32_, *Al S*_b_ (*S*_20_), and potentially *Al S*_36_ (**Supplementary File 5**) have been embedded within class A2/IV. Notably, the different binding mode observed in the complex models of *Ah S*_28_, *Al S*_6_, *Al S*_14_, *Al S*_18_, and *Al S*_a_ correlate with the phylogenetic or dominance class, class B/II or A3/III, for SRK in *Arabidopsis*^*93*^. These complex models exhibited large negative free energy changes, except for *Al S*_18_, and are thus considered plausible (**Figure 5A and Fig. S12**). Furthermore, considering that both class-I and class-II *Brassica* SRK alleles are embedded within the class A2/IV in *Arabidopsis*, a correlation may exist between the eSRK–SP11 binding modes and the phylogenetic/dominance classes in Brassicaceae. However, owing to the limited availability of paired *Arabidopsis* SRK/SP11 sequences in the sequence database, we were unable to investigate this correlation in depth. Nevertheless, performing protein crystallization for the proteins associated with these allelic classes may be crucial to gain further insights into the evolution–structure relationship.

We have thus demonstrated that experimentally curated homologous sequences and additional manipulations to align the Cys residues yielded more plausible protein models than those predicted using the original AlphaFold2. Meanwhile, the improvement in structure prediction for eSRK–SP11 complexes using paired MSA was largely attributed to the architecture of AlphaFold2/ColabFold to consider the coevolutionary signals embedded in the MSA. This implies that as more annotated sequence pairs of eSRK and SP11 become available in the future, their prediction accuracy will enhance. We anticipate that a collaborative effort between experimental and computational biology will unravel the complexities of self-incompatibility in Brassicaceae.

## ASSOCIATED CONTENT

### Data Availability

The A3M-formatted MSA files and the computationally predicted eSRK, SP11, and eSRK–SP11 models using the MSAs are freely accessible at Zenodo, https://doi.org/10.5281/zenodo.8047768.

## AUTHOR INFORMATION

### CRediT authorship contribution statement

**Tomoki Sawa:** Methodology, Investigation, Data Curation, Writing - Original Draft; **Yoshitaka Moriwaki:** Conceptualization, Methodology, Investigation, Validation, Data Curation, Writing - Review & Editing, Visualization, Supervision, Project administration, Funding acquisition; **Hanting Jiang:** Investigation; **Kohji Murase:** Conceptualization, Resources, Writing - Review & Editing; **Seiji Takayama:** Writing - Review & Editing, Supervision; **Kentaro Shimizu:** Supervision; **Tohru Terada:** Writing - Review & Editing, Supervision, Funding acquisition.

### Notes

The authors declare that they have no known competing financial interests or personal relationships that could have appeared to influence the work reported in this paper.

## ACKNOWLEDGMENT

This study was conducted using the TSUBAME3.0 supercomputer at Tokyo Institute of Technology, the General Projects on supercomputer “Flow” at Information Technology Center, Nagoya University, and the FUJITSU Supercomputer PRIMEHPC FX1000 and FUJITSU Server PRIMERGY GX2570 (Wisteria/BDEC-01) at the Information Technology Center, The University of Tokyo. We would like to thank Editage (www.editage.jp) for English language editing.

## ABBREVIATIONS

SRK: *S* receptor kinase
SP11: *S*-locus protein 11
SI: self-incompatibility
pLDDT: predicted local distance difference test
MSA: multiple sequence alignment.

---

**Figure S1.**
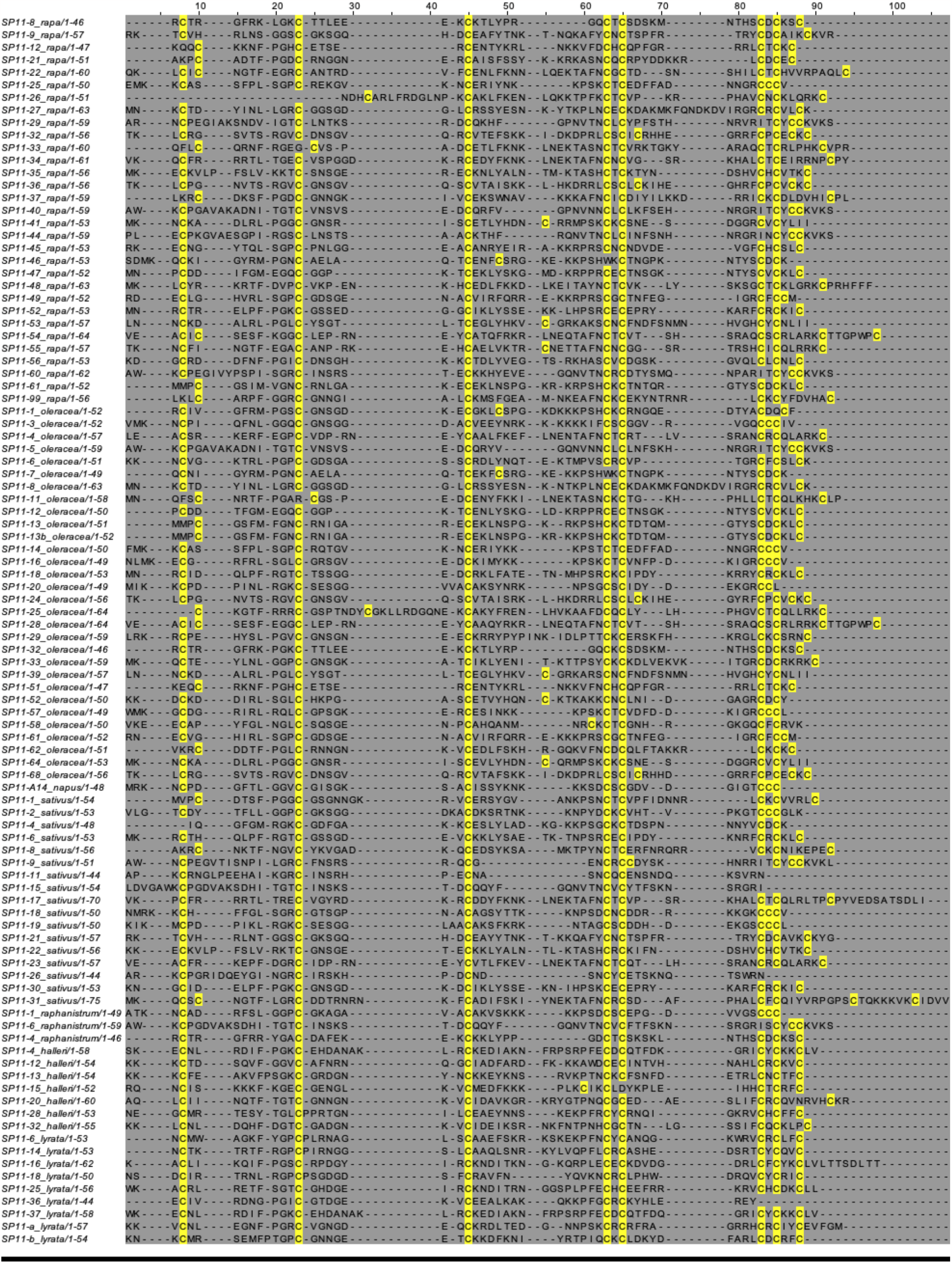
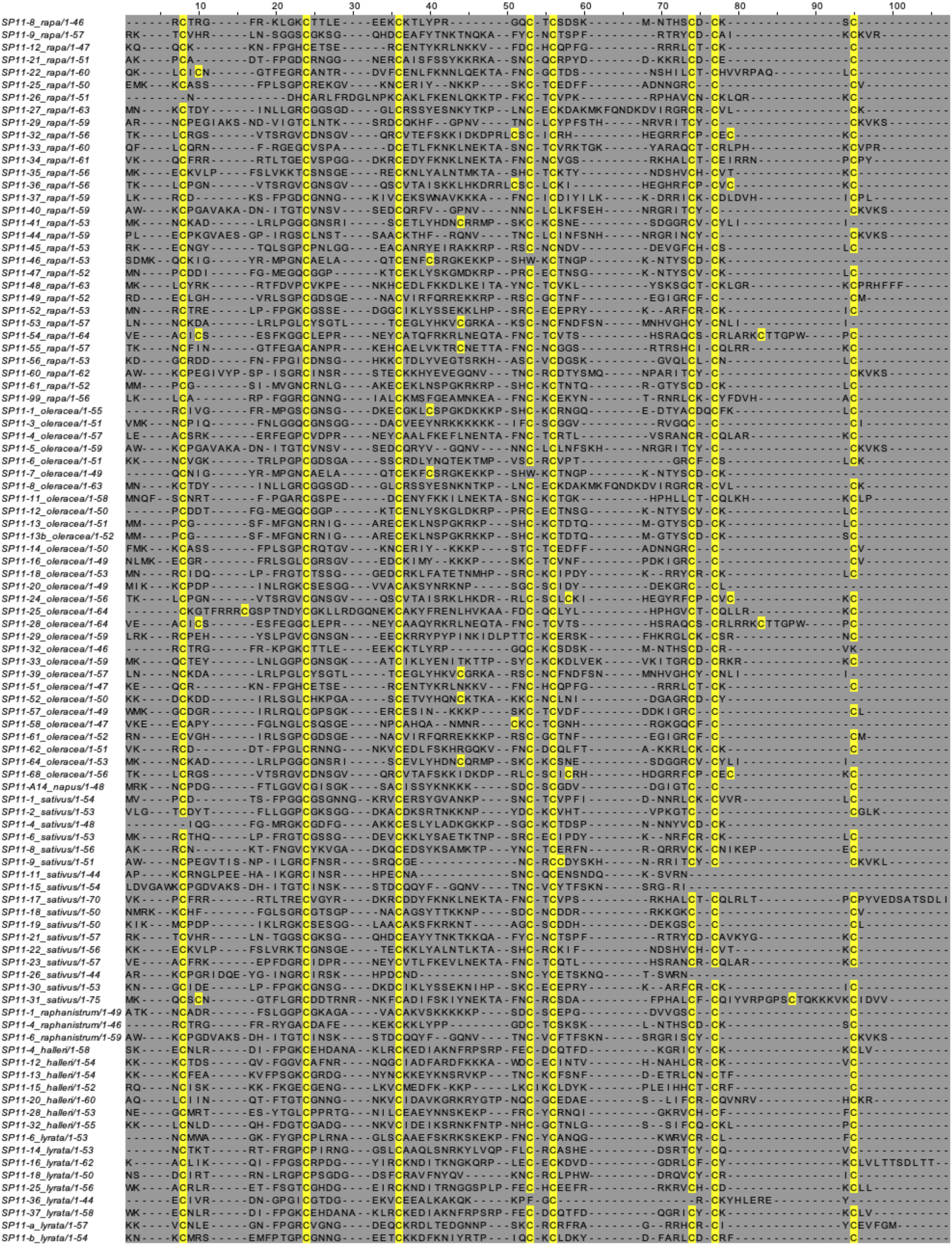
“seq_MSA” (upper) and “cys-seq MSA” (bottom). Cysteine residues are highlighted in yellow. This alignment has been visualized using Jalview 2.11.2.6.

**Figure S2.**
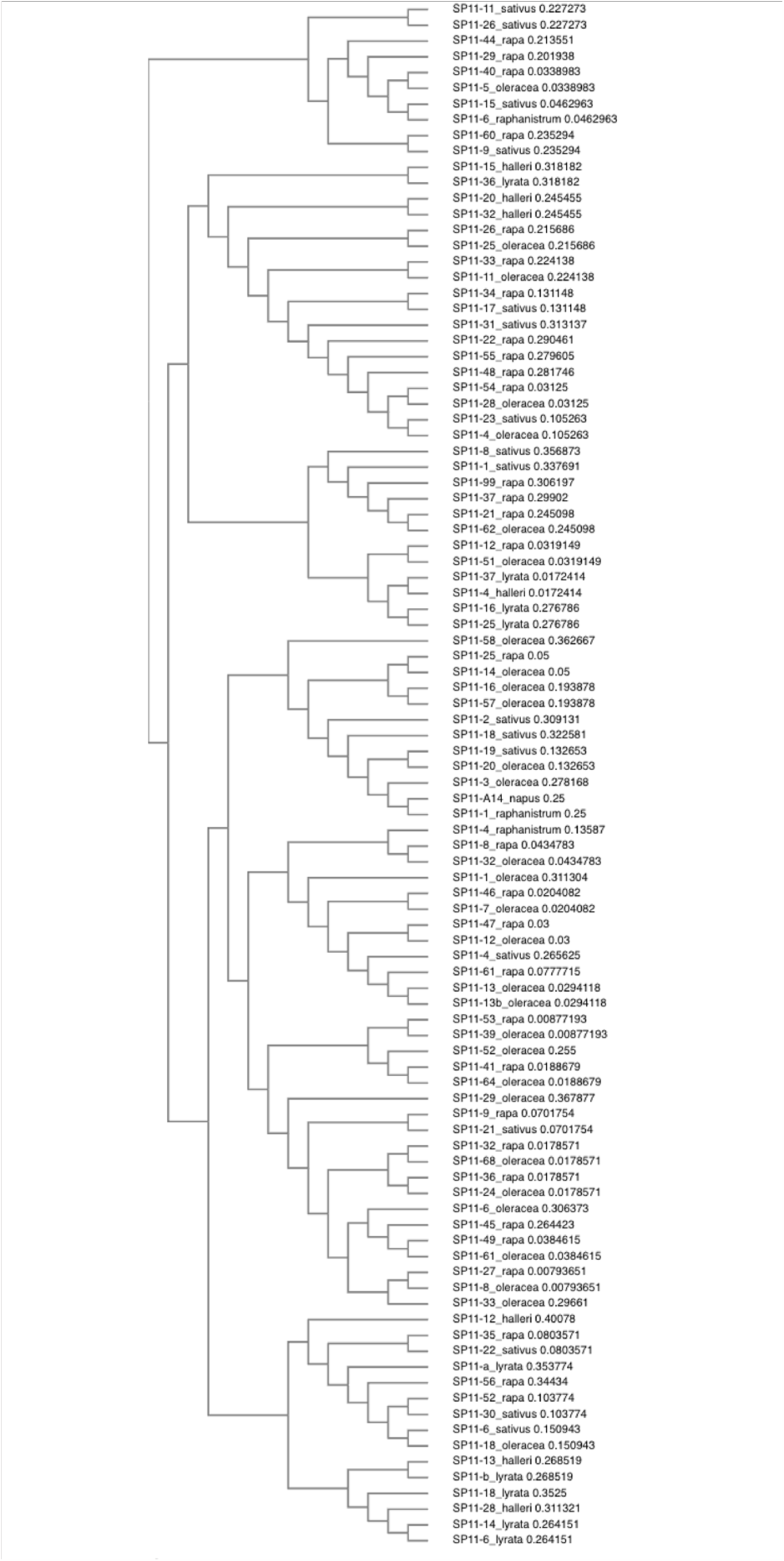
Phylogenetic tree of the 98 SP11 sequences. This figure has been created using Clustal Omega 1.2.4.

**Figure S3.**
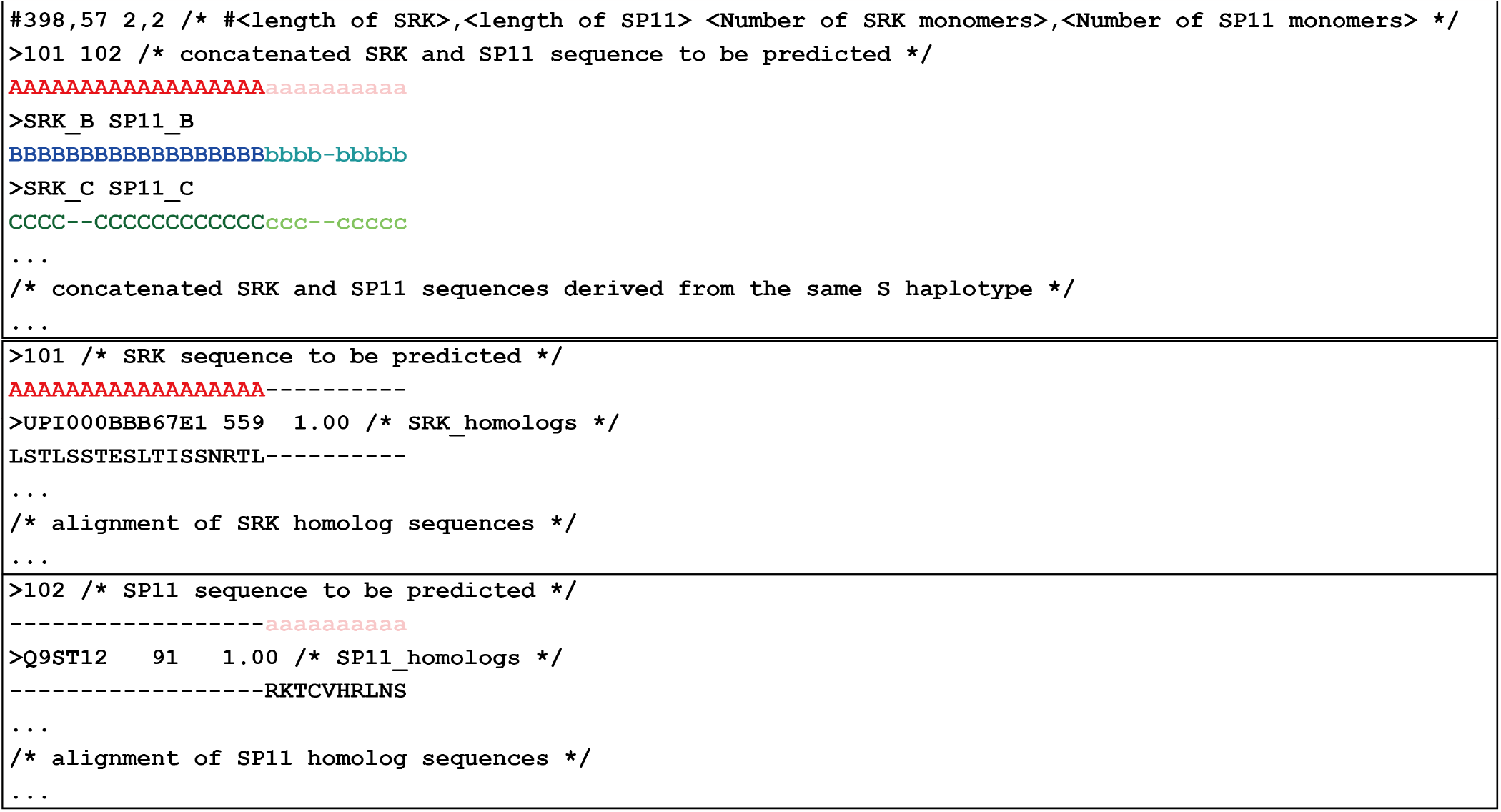
Sample a3m file for structure prediction of cognate SRK and SP11 complex. The amino acid length of SRK and SP11, and the number of SRK and SP11 monomers are specified in the first line starting with # character (e.g. 398,57 2,2). Concatenated SRK and SP11 sequence to be predicted are placed in the first row of the first block, followed by the remaining 97 concatenated SRK–SP11 sequence pairs. SRK and SP11 homolog sequence alignments with gaps are placed in the second and third blocks, respectively.

**Figure S4.**
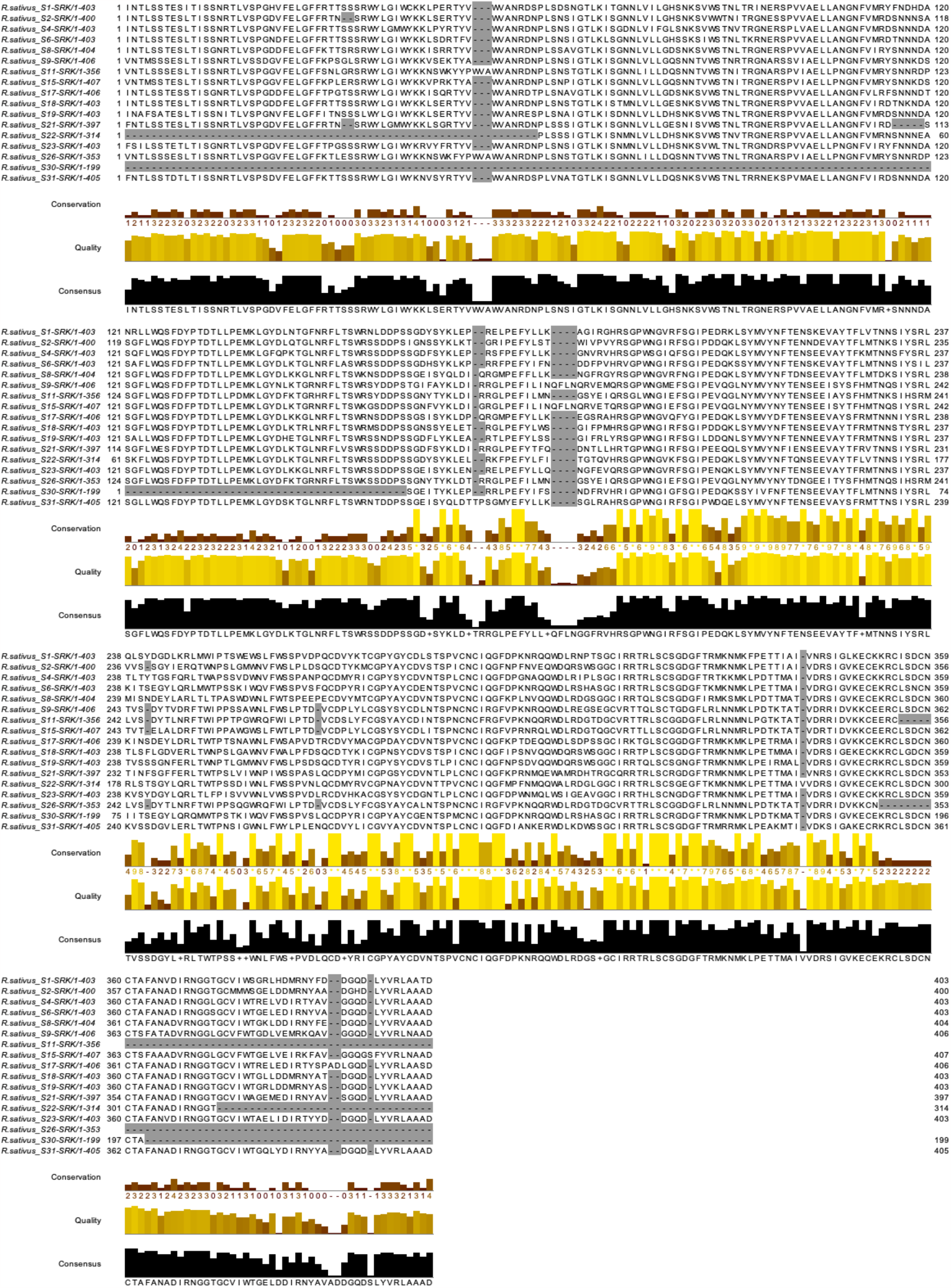
Sequence alignment of *Rs* SRKs. Sequence regions used to run the structure prediction are shown. This alignment has been visualized using Jalview 2.11.2.6.

**Figure S5.**
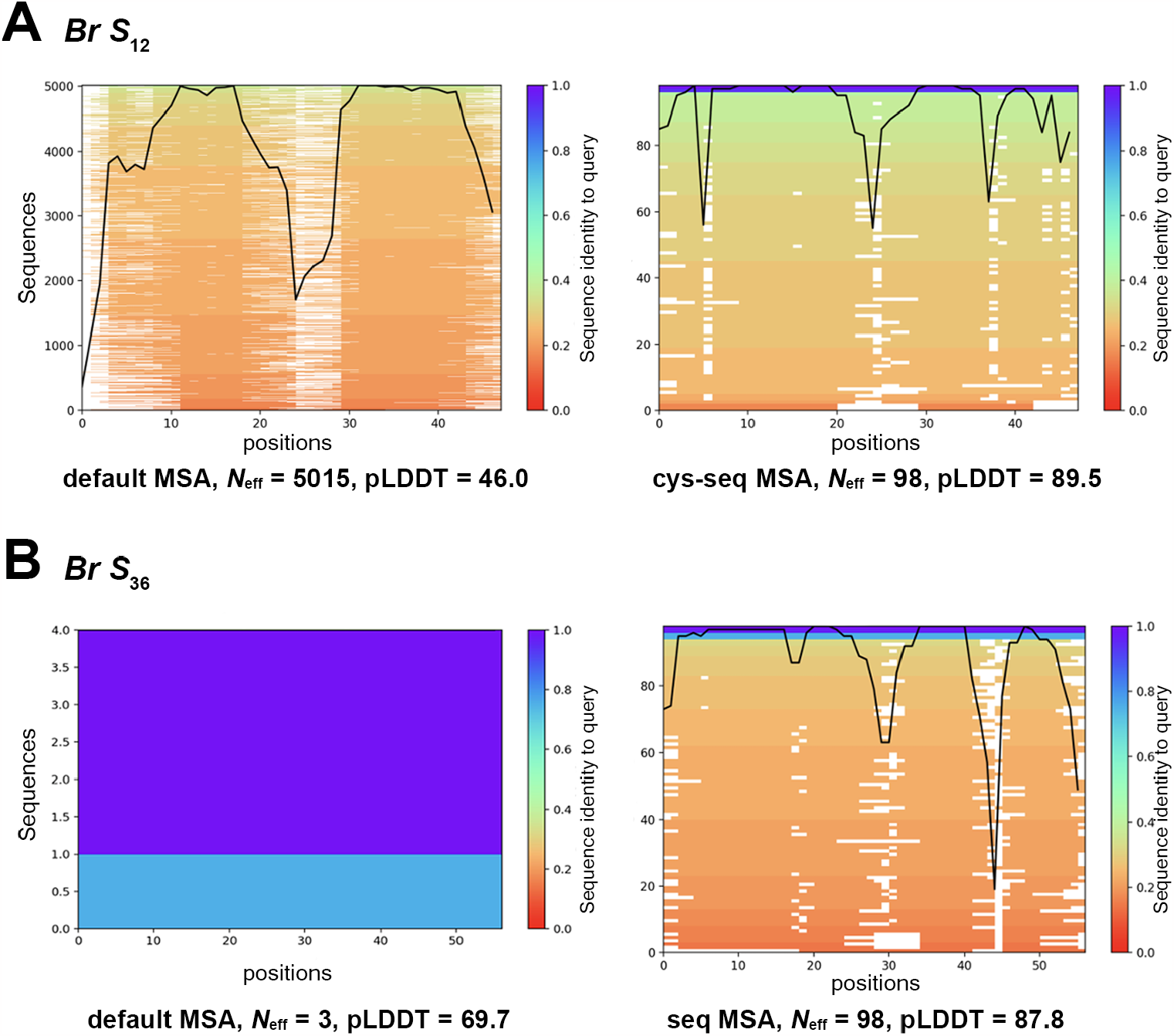
MSA coverage used for structural prediction. The diagrams depict the range of amino acid sequences comprising the input MSA file (in a3m format) that was used to predict the structure of (A) *Br S*_12_-SP11 and (B) *Br S*_36_-SP11. The default MSA (uniref30_2202 + bfd_mgy_colabfold database) and the best MSA (cys-seq for *Br S*_12_ and seq MSA for *Br S*_36_) are shown in the left and right panels, respectively. Each amino acid sequence consisting of the MSA is colored according to its identity to the query sequence. The black line illustrates the ratio of present amino acids (not gaps) relative to the amino acid positions in the query sequence. The diagrams were automatically generated by (Local)ColabFold.

**Figure S6.**
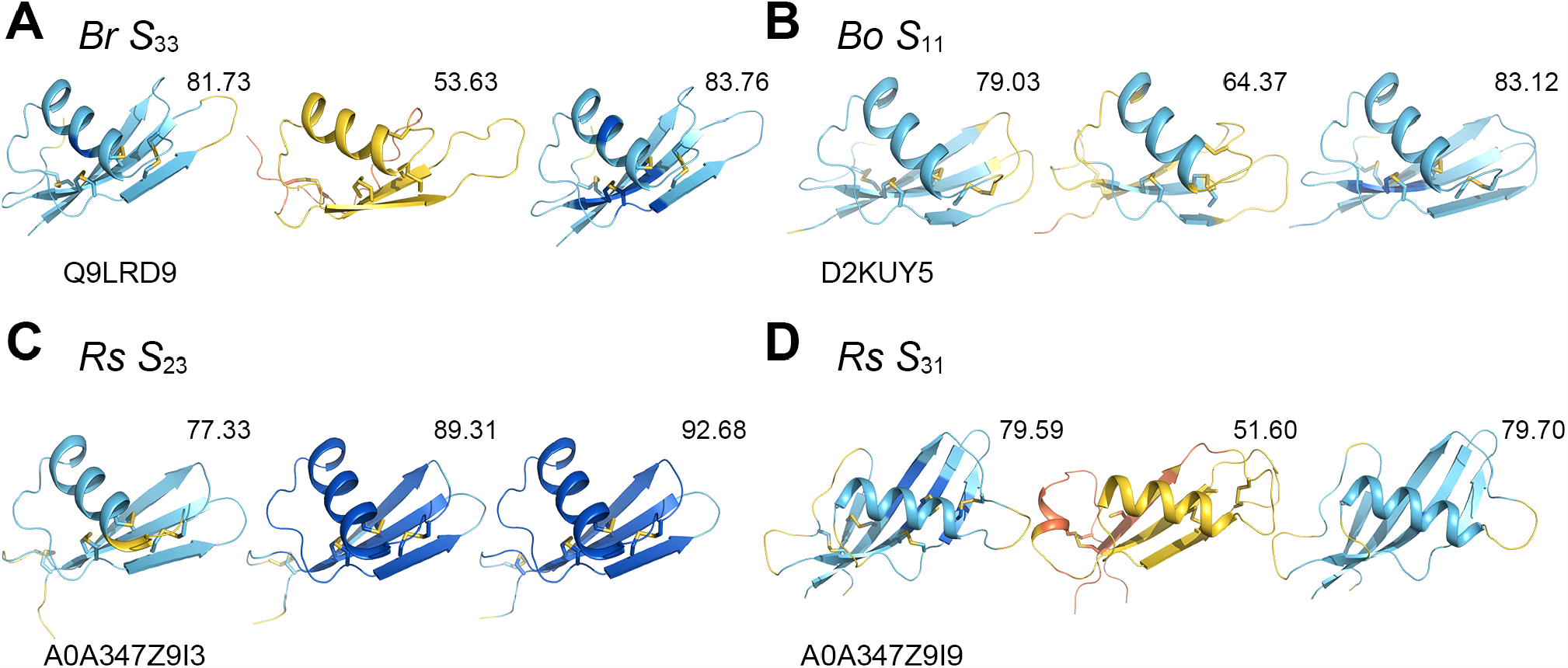
Improved SP11 models with custom MSAs. (A–D) Predicted SP11 structures of *Br S*_33_ (A), *Bo S*_11_ (B), *Rs S*_23_ (C), and *Rs S*_31_ (D) haplotypes. The predicted model deposited in AlphaFold DB (version 2022-11-01), one with the default MSA, and the best one among the four custom MSAs are depicted in the left, middle, and right panels, respectively. UniProt accession IDs are shown under the model of AlphaFold DB. The mean per-residue pLDDT value is shown on the upper right corner of each model. The structures are colored according to the AlphaFold pLDDT model confidence.

**Figure S7.**
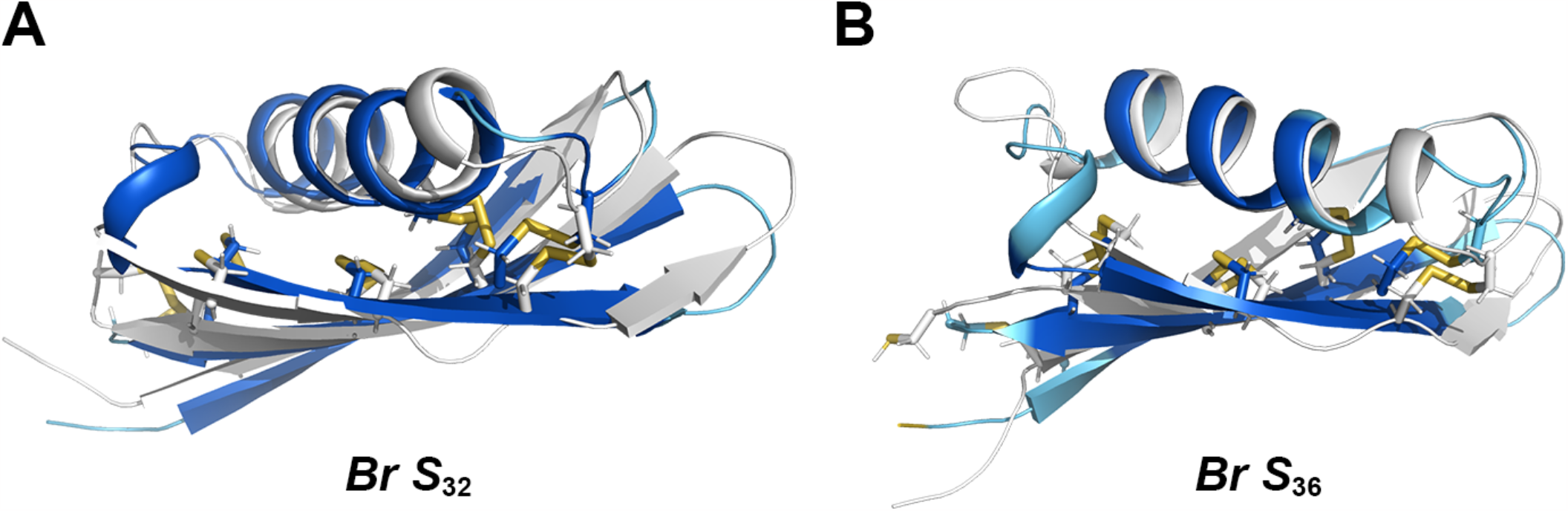
Superposition of the SP11 models predicted in our previous and current study. Panels A and B show the predicted models of *Br S*_32_- and *S*_36_-SP11, respectively. The best models predicted by ColabFold in this study are colored using the pLDDT values. The models from our previous study are shown in white.

**Figure S8.**
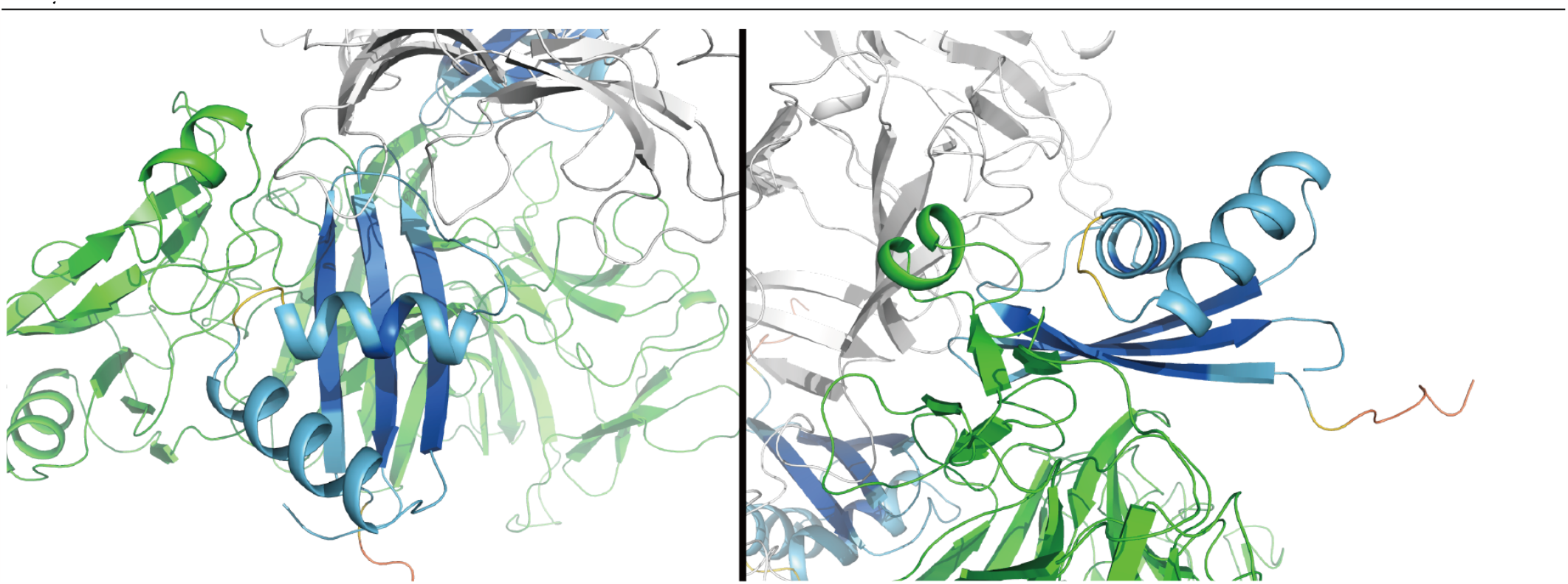
A predicted model of *Br S*_26_-SRK–SP11. Close-up view of the interface where the two SRK molecules (white and green) and one SP11 molecule make contact. The second helix of SP11 does not make contact with the SRK molecules.

**Figure S9.**
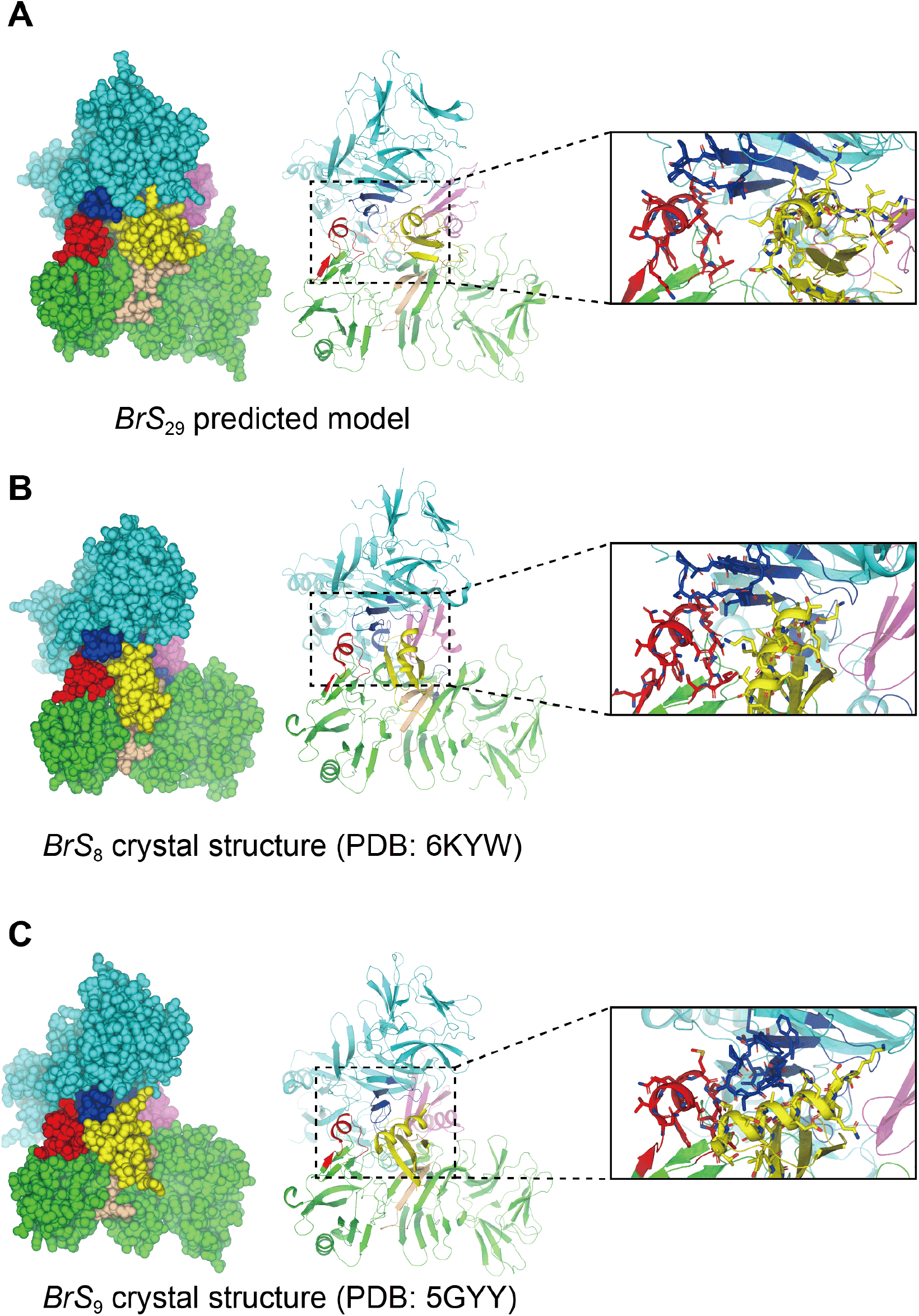
Structural comparison of the binding interface between HV-III of eSRK and SP11. (A) A predicted model of *Br S*_29_ as a representative of the class-II eSRK–SP11 complexes. (B) The crystal structure of *Br S*_8_ (PDB: 6KYW). (C) The crystal structure of *Br S*_9_ (PDB: 5GYY). The two eSRK monomers have been colored in green and cyan and the two SP11s in yellow and purple. In this figure, the position and orientation of eSRK chain A (green) are aligned. The HV regions I, II, and III of eSRK are colored in red, deep blue, and light brown, respectively. HV-I and III belong to the same eSRK chain, whereas HV-II belongs to the other chain. Sphere and cartoon representations are shown in the left and center panels, respectively. A close-up view of the binding interface between eSRK and SP11 are shown in the right panel.

**Figure S10.**
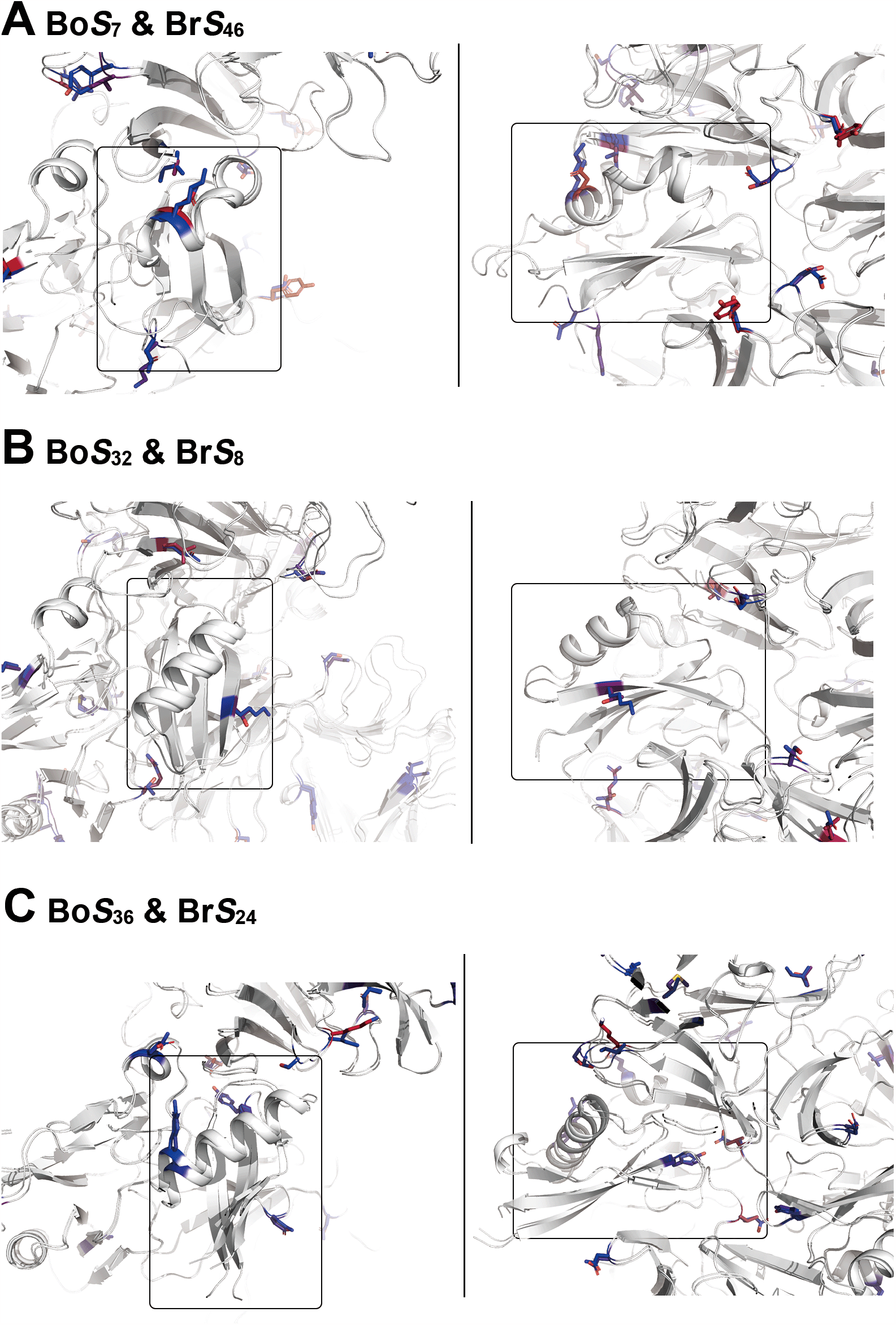
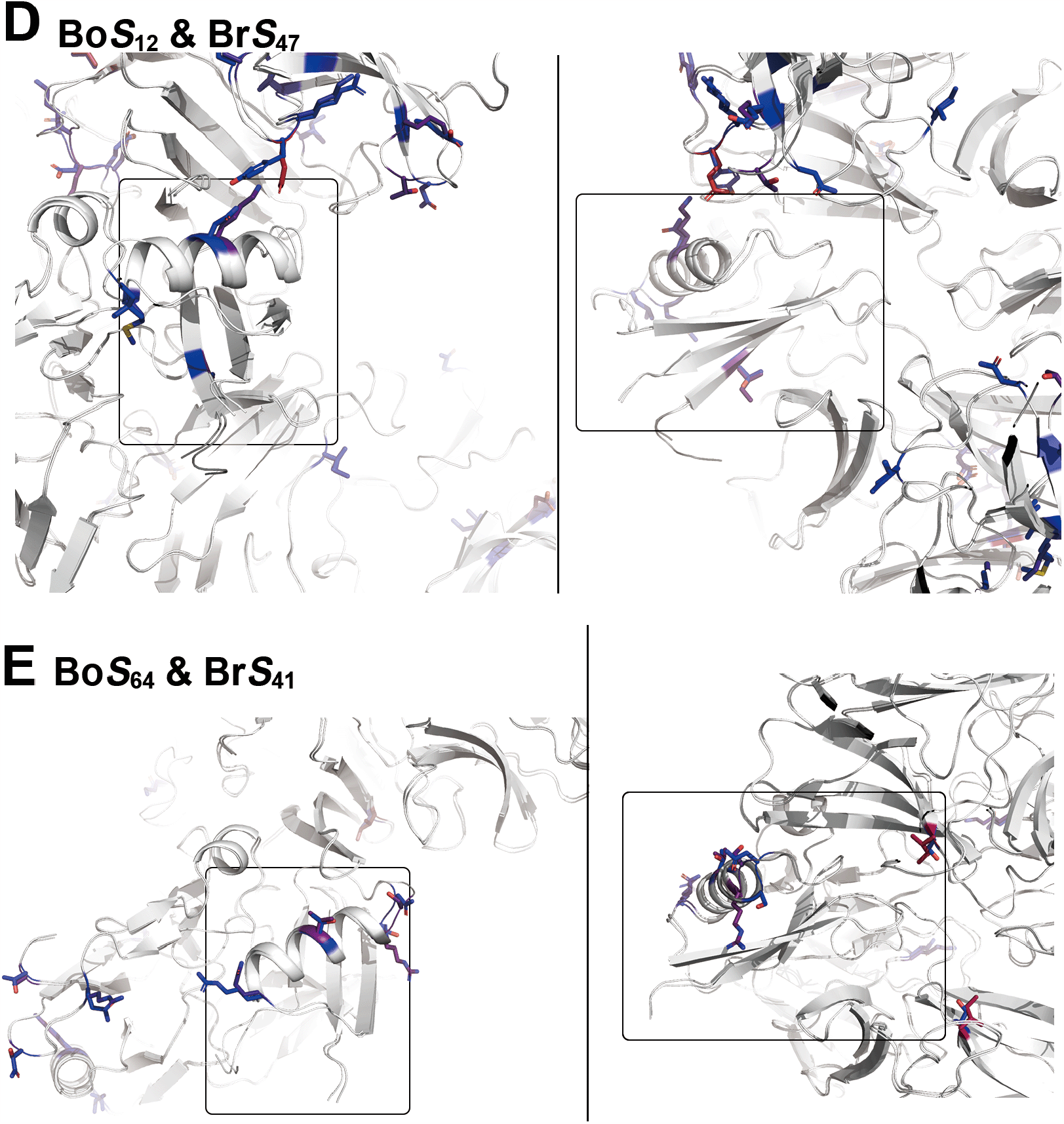
Superposition of the predicted eSRK–SP11 complex models that have highly similar sequence identity. (A) *Bo S*_7_ and *Br S*_46_, (B) *Bo S*_32_ and *Br S*_8_, (C) *Bo S*_36_ and *Br S*_24_, (D) *Bo S*_12_ and *Br S*_47_, and (E) *Bo S*_64_ and *Br S*_41_. Mutated residues are shown as sticks and in color according to their difference in the BLOSUM90 matrix. The position of SP11 is highlighted inside a black rectangle.

**Figure S11.**
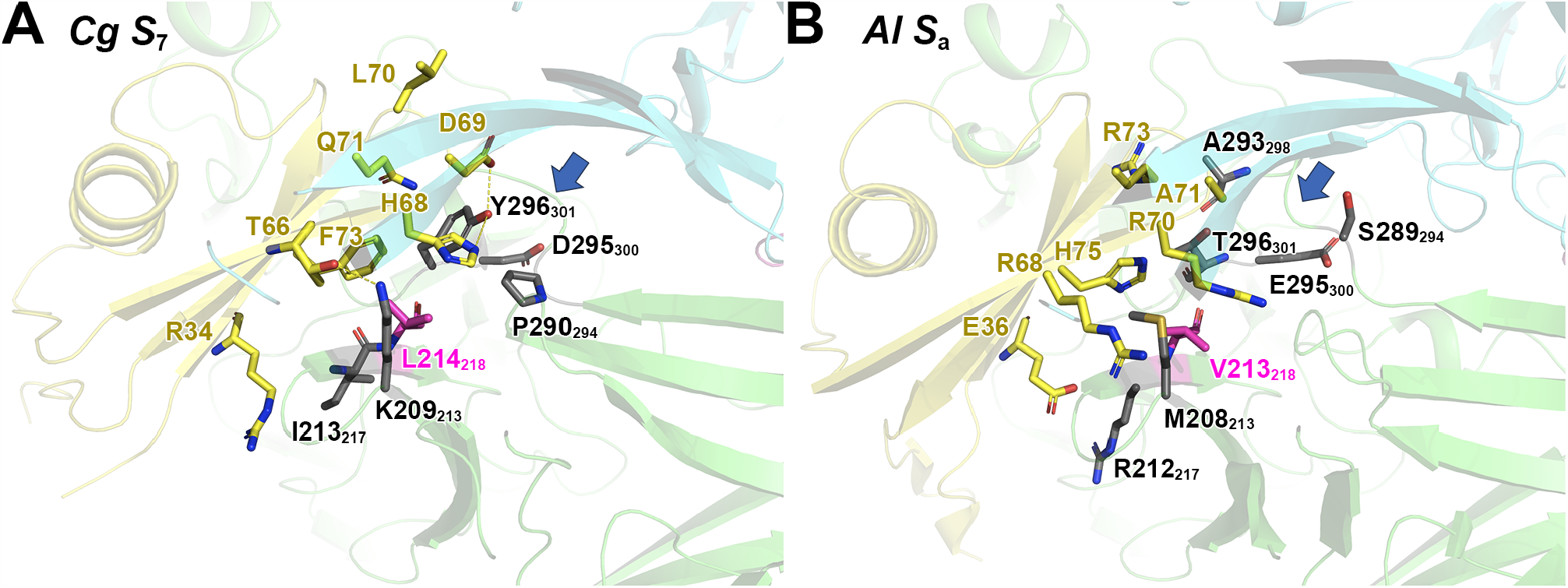
Predicted *Cg S*_7_ and *Al S*_a_ complex models. Close-up views of the eSRK–SP11 interface of *Cg S*_7_ (A) and *Al S*_a_ (B). The two eSRK monomers have been colored in green and cyan and SP11 in yellow. The subscripts represent the residue number in the eSRKa(7)a chimera protein. The arrow indicates a loop structure corresponding to residues 294–300 in eSRKa(7)a.

**Figure S12.**
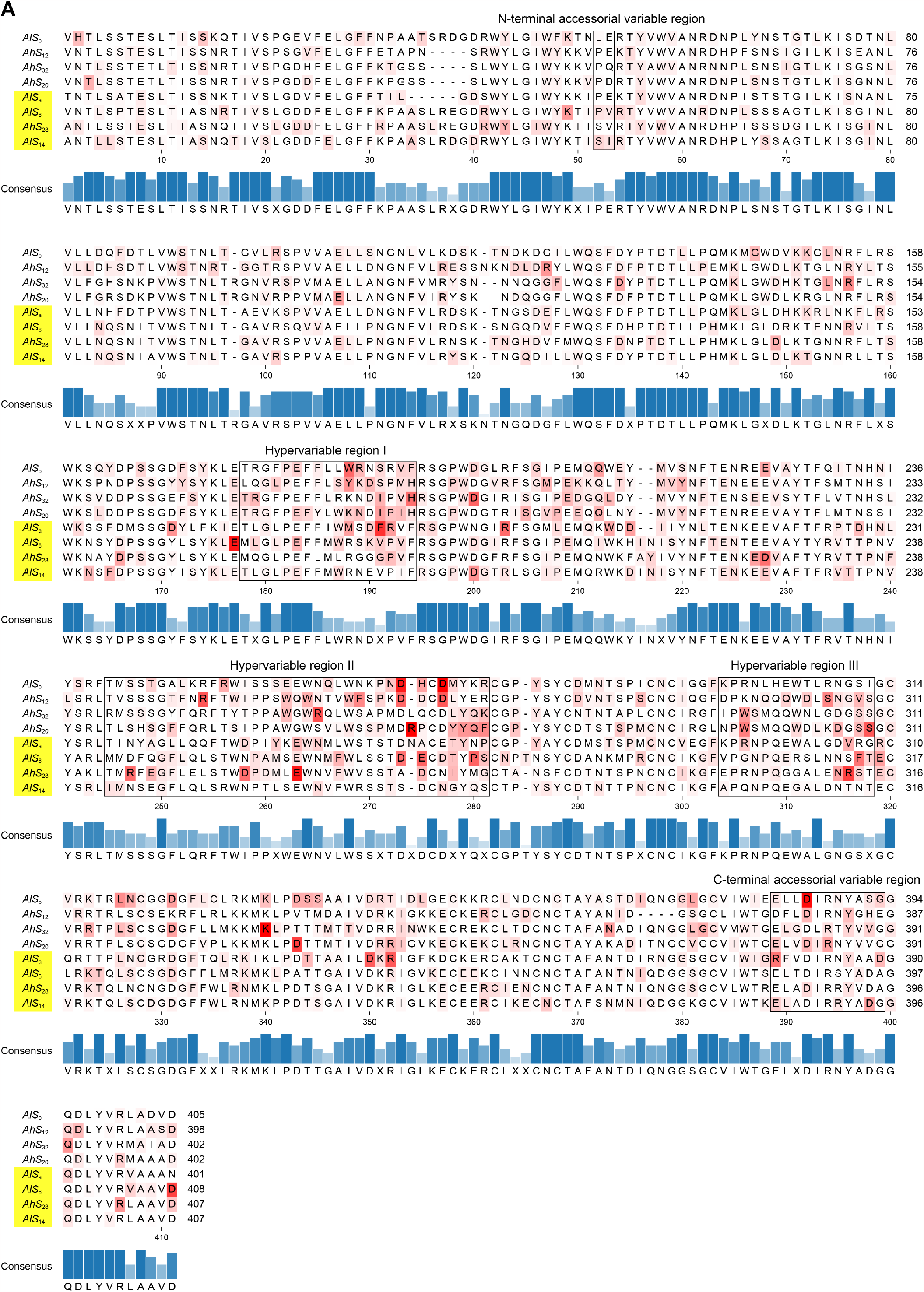
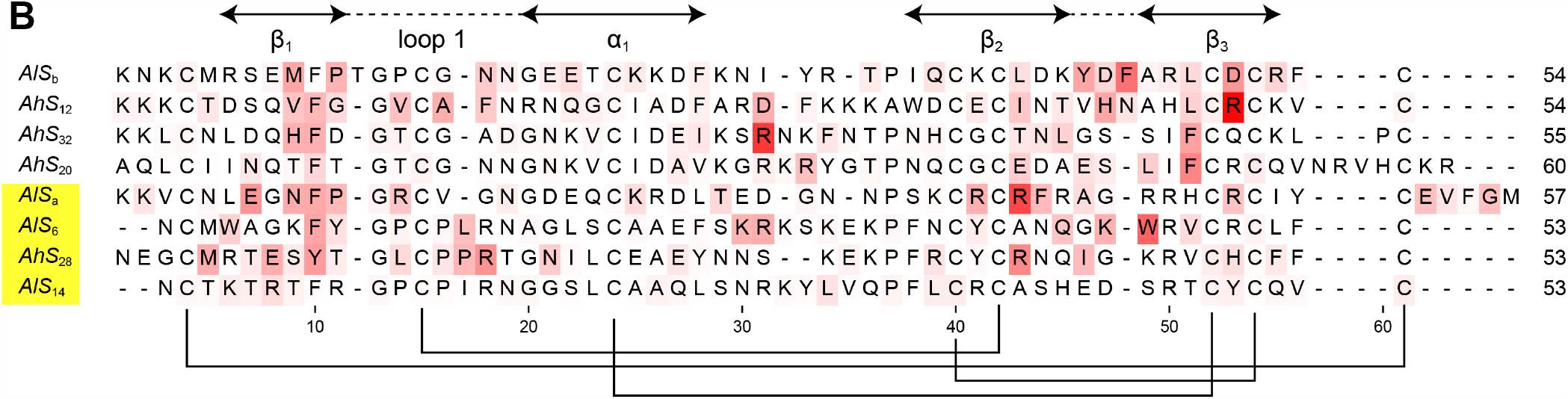
Per-residue energy decomposition analysis for the predicted *Al*/*Ah* eSRK–SP11 complex models. (A–B) Sequence alignment of *Al* and *Ah* SRKs (A) and SP11s (B). Haplotypes predicted to have the different binding mode from *Br S*_8_/*S*_9_ are highlighted in yellow. The variable regions are indicated by black borders.

**Table S1.**
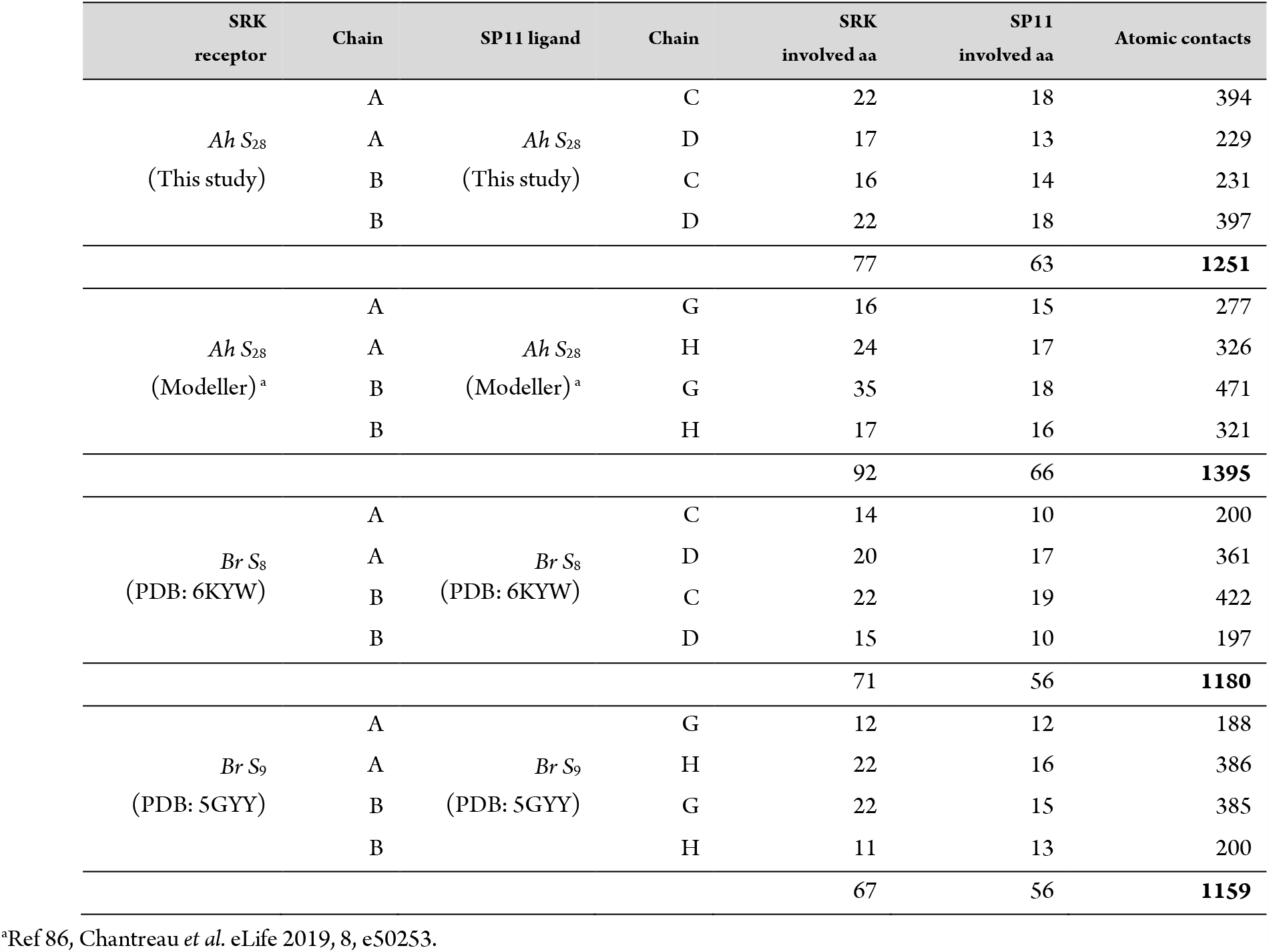
The number of atomic contacts between the eSRK and SP11 molecules.

An example to calculate the number of atomic contacts and involved residues on PyMOL. The get_raw_distance command is available in Psico module (https://pymolwiki.org/index.php/Psico).

~~~
delete everything
fetch 5gyy # load PDB: 5GYY, Br S9 SRK-SP11 crystal structure
select not polymer # select non-polypeptide residues
remove sele # delete non-polypeptide residues
delete dist* # delete existing distance objects
# make distance objects between chain A and G with a 5.0 Å atom-atom cut-off
distance dist1, chain A, chain G, cutoff=5
dist_obj1 = cmd.get_raw_distances(“dist1”)
print(len(dist_obj1)) # 188
num_residues1 = len(set([i[0][1] for i in dist_obj1]))
num_residues2 = len(set([i[1][1] for i in dist_obj1]))
print(num_residues1) # 12
print(num_residues2) # 12
distance dist2, chain A, chain H, cutoff=5
dist_obj2 = cmd.get_raw_distances(“dist2”)
print(len(dist_obj2)) # 386
num_residues1 = len(set([i[0][1] for i in dist_obj2]))
num_residues2 = len(set([i[1][1] for i in dist_obj2]))
print(num_residues1) # 22
print(num_residues2) # 16
distance dist3, chain B, chain G, cutoff=5
dist_obj3 = cmd.get_raw_distances(“dist3”)
print(len(dist_obj3)) # 385
num_residues1 = len(set([i[0][1] for i in dist_obj3]))
num_residues2 = len(set([i[1][1] for i in dist_obj3]))
print(num_residues1) # 22
print(num_residues2) # 15
distance dist4, chain B, chain H, cutoff=5
dist_obj4 = cmd.get_raw_distances(“dist4”)
print(len(dist_obj4)) # 200
num_residues1 = len(set([i[0][1] for i in dist_obj4]))
num_residues2 = len(set([i[1][1] for i in dist_obj4]))
print(num_residues1) # 11
print(num_residues2) # 13
print(len(dist_obj1) + len(dist_obj2) + len(dist_obj3) + len(dist_obj4)) # 1159
~~~

