## Supplementary figures and images for "Comprehensive computational analysis of the molecular mechanism of self-incompatibility in Brassicaceae using improved structure prediction"

### Supplementary File 3

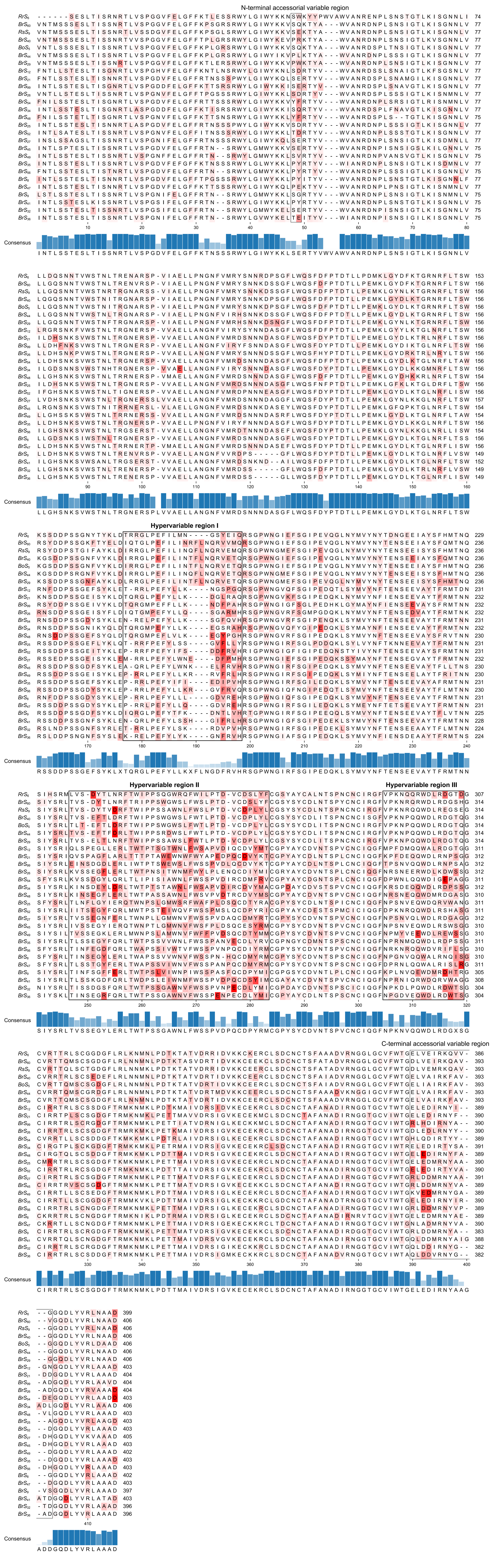

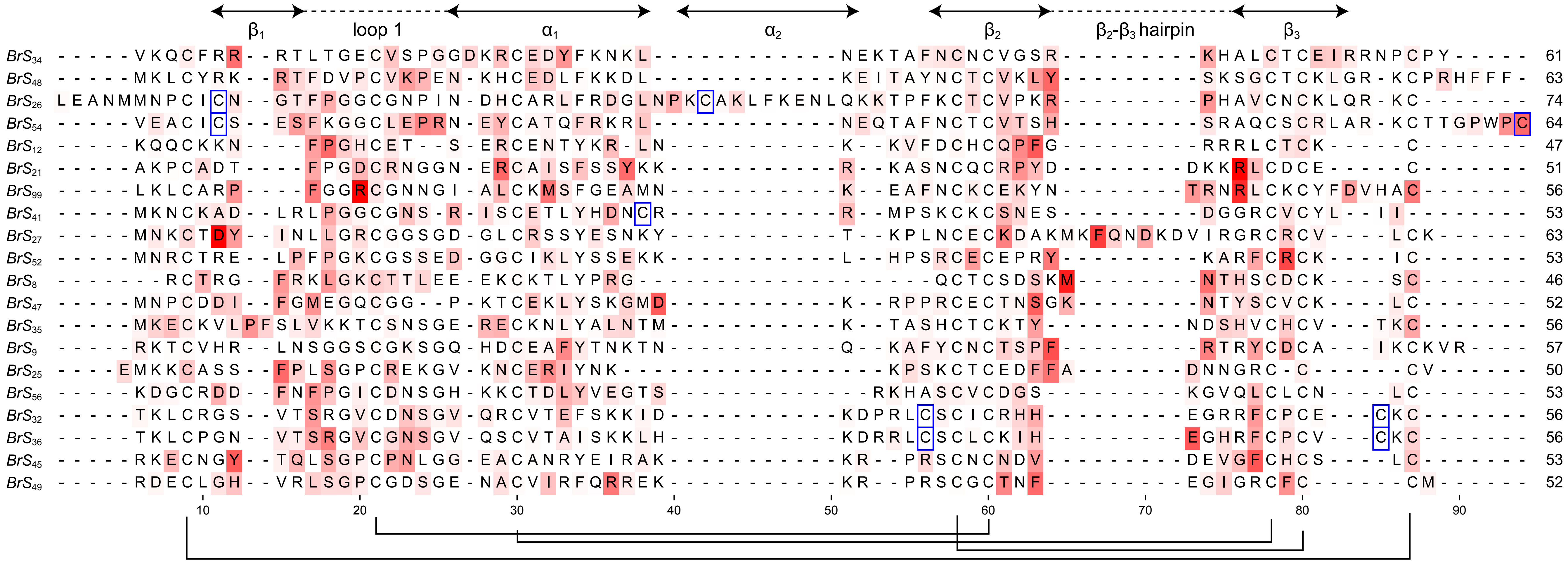

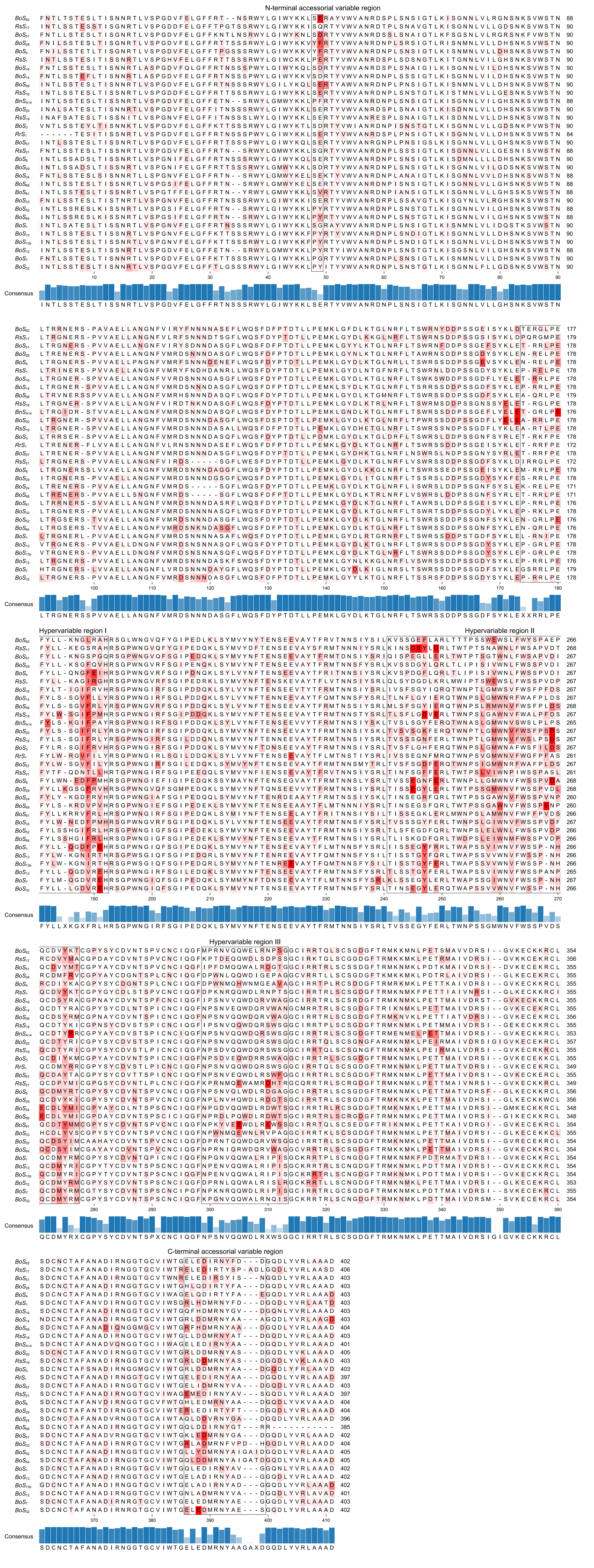

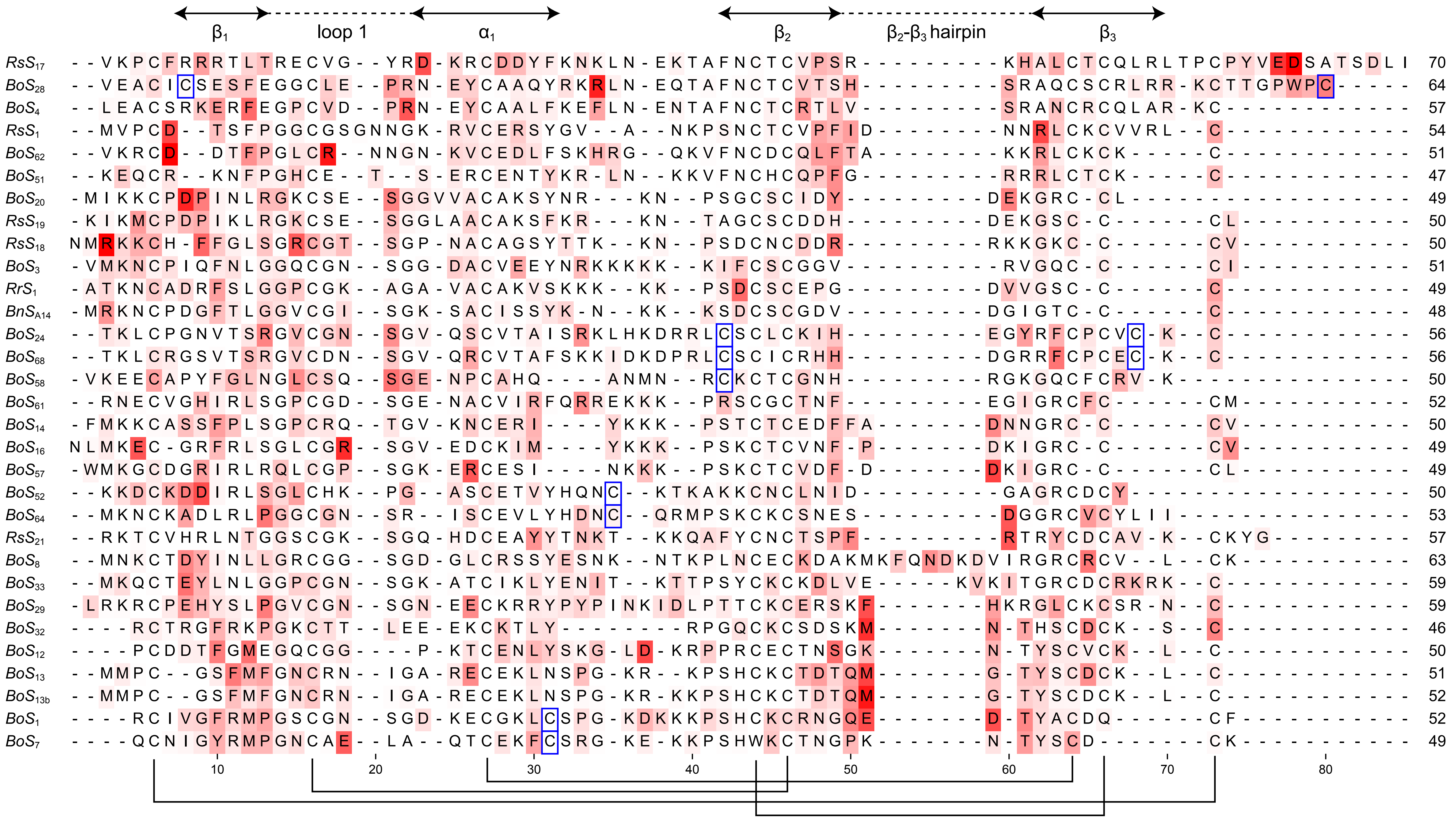

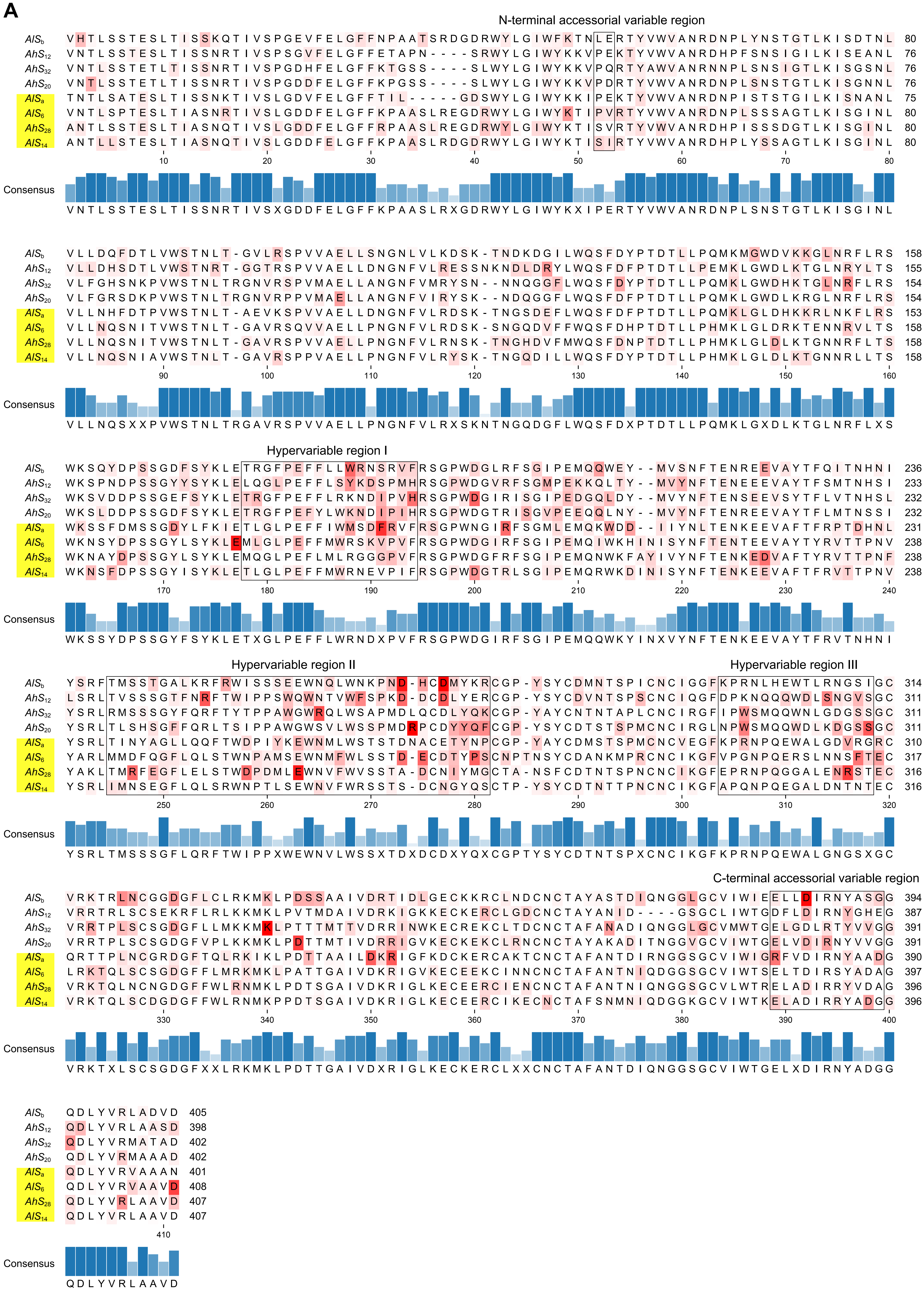

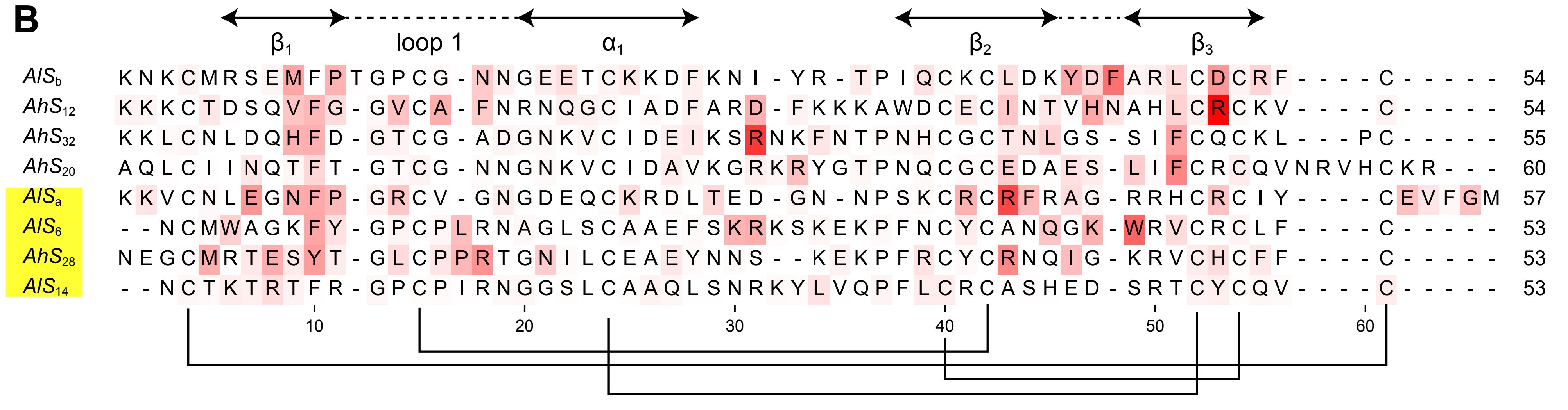
