## Supplementary File 5 for "Comprehensive computational analysis of the molecular mechanism of self-incompatibility in Brassicaceae using improved structure prediction"

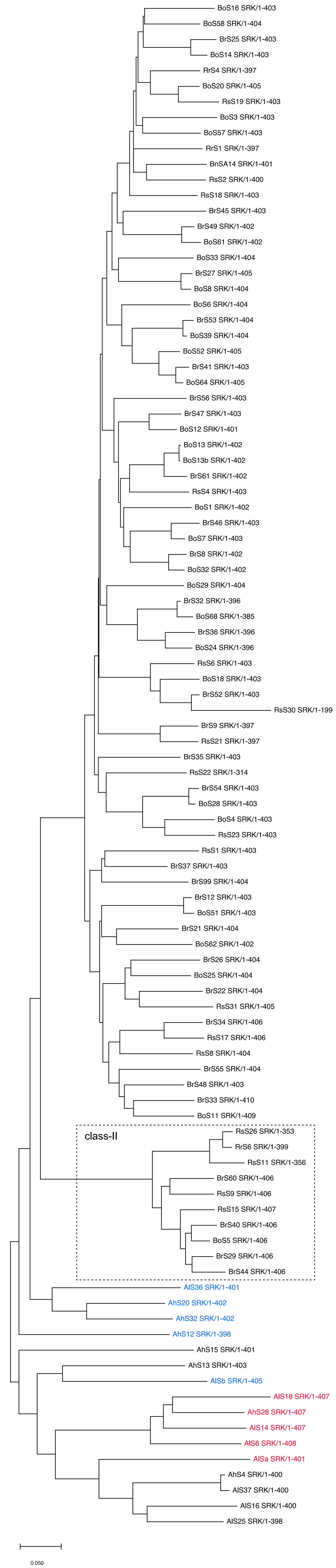


*Haplotypes of *Ah* and *Al* showing a similar binding mode to the crystal structure of *Br* $S_8$ or $S_9$ are colored in blue (Figure 5B) , and those with the different binding mode (Figure 5A) are in red.
